# The role of folate receptor α in the partial rejuvenation of dentate gyrus cells. Improvement of cognitive function in elderly mice

**DOI:** 10.1101/2023.02.01.526619

**Authors:** A Antón-Fernández, R Cuadros, R Peinado-Cahuchola, F Hernández, J Avila

## Abstract

In this work, we have studied the effect of small compounds in the partial rejuvenation of dentate gyrus cells by measuring the improvement of cognitive functions in elderly mice.

Aging has been related to a change in DNA methylation and some one-carbon metabolites linked to that methylation process, like vitamin B12, folate or methionine have been involved in cognitive performance during aging. However, their role in this process and the possible mechanisms behind its cognitive effects are still unclear. Through direct infusion of these molecules in the dentate gyrus we have tested their effects on cognition in elderly mice. Only positive results were found for folate. A partial rejuvenation of dentate gyrus cells related to an increase in neuroplasticity by reorganizing extracellular matrix structures and rising the expression of juvenile genes like GluN2B was found. Since folate is involved in several cellular pathways in addition to DNA methylation, we have focused in its interaction with its folate receptor alpha (FRα), a protein that is present at the cell nucleus, acting as transcription factor. We have found that most of folate effects on brain would be mediated by the activation of FRα. In addition, we propose that the mechanism for cell rejuvenation by folate, or other FRα binding molecules, may involve the expression of proteins, like SOX2, a Yamanaka factor present in young neurons. Thus, the use of molecules that activate the FRα pathway could constitute an interesting strategy to be considered for the study of brain rejuvenation.

## 1. Introduction

Epigenetic changes are hallmark features of aging (Lopez-Otin et al., 2013). Indeed, organismal aging can be monitored by studying several epigenetic chromatin markers (Sen et al., 2016). Given that aging-dependent epigenetic changes can be modified to younger levels, the reversion of some aging characteristics is feasible (Benayoun et al., 2015; Pollina and Brunet, 2011; Zhang et al., 2015). Nowadays that process is possible without loss of cellular identity in a procedure called partial reprogramming (Olova et al., 2019). This partial reprogramming has been achieved as a way to rejuvenate old cells without loss of somatic identity (Olova et al., 2019). The partial reprogramming can be achieved by the cycle expression of the so-called Yamanaka Factors (YF) (Takahashi and Yamanaka, 2006; Watanabe et al., 2013). Under that expression, there is an amelioration of some hallmarks of aging in peripheral tissues (Chondronasiou et al., 2022; Ocampo et al., 2016; Sarkar et al., 2020). It should be quoted that induced pluripotent stem cells (iPSCs) can be derived from several, but not all, types of somatic cells. Postmitotic neurons are not able to reprogrammate into induced pluripotent stem cells (iPSCs) by expressing only YF. For that reprogrammation would be necessary to induce cell proliferation in addition to YF expression (Kim et al., 2011). However, YF can promote partial reprogramming (rejuvenation) in neuronal cells from old mice, facilitating a cognitive improvement (Rodriguez-Matellan et al., 2020).

Here we tested a strategy to rejuvenate aging brain cells through specific partial reprogramming (Chen and Skutella, 2022) but using small simple compounds (Hou et al., 2013) instead of YF. In this regard, YF may reprogram neurons in the brains of old mice through the following: a) an increase in adult hippocampal neurogenesis (AHN)(Rodriguez-Matellan et al., 2020); b) epigenetic changes (methylation) in histones or DNA (Planello et al., 2014; Rodriguez-Matellan et al., 2020) ; or c) acting YF as transcription factors (Yamanaka, 2008). Thus, here we tested whether small compounds could replicate the effects of YF. Our findings showed that the compounds tested had no significant impact on AHN. On the second point, it is known that YF control epigenetic changes like those related to the regulation of histone and DNA methylation (Ocampo et al., 2016; Rodriguez-Matellan et al., 2020). Changes in the latter sustain the development of the epigenetic clock, which may predict biological (chronological) age (Horvath and Raj, 2018). Histone and DNA methylation involves methyl donors, methyltransferases, and, sometimes, coenzymes like vitamin B12 (Ducker and Rabinowitz, 2017). The universal methyl-donor is methionine but other factors like folate could be involved indirectly (Ducker and Rabinowitz, 2017). It has been reported that YF and their related Thompson factors (Planello et al., 2014) increase the activity of DNA methyltransferases like the DNA methyltransferase (DNMT) family (Lopez-Bertoni et al., 2015; Tsai et al., 2012; Wu et al., 2018). Given that DNA methylation can increase in the presence of methyltransferases, through the presence of methyl-donors like the universal methyl-donor (methionine), folate, or cofactors like vitamin B12 (Ducker and Rabinowitz, 2017), here we studied whether the presence of any of these small compounds could replicate the effects of YF in the amelioration of aging features in the dentate gyrus (DG) (Rodriguez-Matellan et al., 2020). Our results suggest that the presence of folate is enough to achieve amelioration and also to induce changes in the degree of DNA methylation. Furthermore, folate could act on cells by binding to folate receptors, being folate receptor α (FRα) the most studied receptor, which, in addition, could serve as a transcriptional factor (Boshnjaku et al., 2012). Here we tested the capacity of a synthetic FRα-binding peptide (Hulin-Curtis et al., 2020) to mimic folate as an inducer of FRα transcriptional function. Indeed, the brain injection of this peptide resulted in cognitive improvement similar to those achieved with folate injections. Given these observations, this FRα-binding peptide emerges as a potential alternative treatment to folate to improve cognition.

## 2. Methods

### 2.1. Animals

A total of 57 C57BL/6 wild-type male and female mice were bred in the animal facility at the *Centro de Biología Molecular Severo Ochoa*. They were housed in a specific pathogen-free facility under standard laboratory conditions following European Community Guidelines (directive 86/609/EEC) and handled in accordance with European and local animal care protocols (PROEX 62/14 and 291/15). They were housed 4–5 per cage with food and water available ad libitum and maintained in a temperature-controlled environment on a 12/12-h light/dark cycle with light onset at 8 a.m.

### 2.2. Experimental design

The mice were divided into experimental groups (average 16 months old for the vitamin B12 experiment and 21 months old for the S-(5′-Adenosyl)-L-methionine chloride (SAM), folate, and peptide experiments). They were randomly injected in the hilus region of both brain hemispheres (2 μl in each one) with vehicle (phosphate buffered saline (PBS) or distilled water) or one of four molecules of interest: vitamin B12 (50 mg/ml, diluted in water; Vet one, ref: 510218; molecular weight, m.w., 1355 g/mol), SAM (0.25 mg/ml diluted in water; Sigma, ref: A7007; m.w. 398,5 g/mol), folic acid (0.25 mg/ml diluted in PBS; Sigma, ref: F7876-10G; m.w., 441 g/mol), or a synthetic FRα-binding peptide (5 mg/ml, diluted in PBS; Abyntec; m.w., 1121 g/mol). The pH of vitamin B12, methionine, and folic acid solution was adjusted with NaOH to ∼7 to avoid negative effects caused by an overly acidic medium. One week after the intracerebral injections, the mice were assessed using behavioral (open field trial) and memory tests (novel object recognition and Y maze test) to determine the potential functional effects of the metabolites on the central nervous system. Two weeks after surgery, the mice were perfused immediately after completing the Y maze test.

### 2.3. Stereotaxic surgery and intrahippocampal microinjection

After anesthesia with isoflurane, the mice were secured in a stereotaxic frame (Kopf). Holes were drilled bilaterally in the skull at the injection sites (one per hemisphere) using a microdrill with a 0.5 mm bit. The following stereotaxic coordinates were used for intrahippocampal injections (from bregma): anterior– posterior -2.0; lateral 1.4; and dorsoventral -2.2. A 33-gauge needle Hamilton syringe coupled to a syringe pump controller mounted on the stereotaxic frame was used to inject 2 μl of vehicle or metabolites at each site. Injections were given at 0.25 μl/min, after which the needle was left in place for an additional 3 min, stopping another 3 minutes in the halfway. After the injections, the skin was sutured and buprenorphine (Sigma, USA) was given subcutaneously, and the animals were allowed to recover for 1 h on a heating pad before returning to the cage. Animals were treated with ibuprofen (Dalsy in drinking water) for the following 5 days. The mice remained in the cage for an additional week before the start of behavior tests and were sacrificed 2 weeks after surgery (**SI 1**).

### 2.4. CldU injection protocol

The mice received three injections every 2 h (200 ul/injection, intraperitoneally in alternative sites of the body) of 5-Chloro-2’-deoxy-Uridine (CldU, Sigma reference C6891; i.p. 42.75 mg/Kg body weight) one week after cranial surgery and microinjections of the solutions and just one week before the sacrifice. This approach allowed us to monitor the effects of the treatments on the proliferation of new cells in the subgranular zone (SGZ), a key region of the brain for neurogenesis, labeling one week-old new generated cells.

### 2.5. Animal sacrifice and tissue processing

The mice were anesthetized with an intraperitoneal pentobarbital injection (Dolethal, 60 mg/kg body weight) and transcardially perfused with saline. Brains were separated into two hemispheres. One hemisphere was removed and fixed in 4% paraformaldehyde in 0.1 M phosphate buffer (PB; pH 7.4) overnight at 4°C. The next day, it was washed three times with 0.1 M PB and cut along the sagittal plane using a vibratome (Leica VT2100S). Serial parasagittal sections (50 mm thick) were cryoprotected in a 30% sucrose solution in PB and stored in ethylene glycol/glycerol at 20°C until they were analyzed. Regarding the other hemisphere, the hippocampus was rapidly dissected on ice and frozen in liquid nitrogen for other studies.

### 2.6. Immunofluorescence techniques

For immunofluorescence experiments, free-floating serial sections (50-μm thick) were first rinsed in PB and then pre-incubated for 2 h in PB with 0.25% Triton-X100 and 3% normal serum of the species in which the secondary antibodies were raised (Normal Goat Serum / Normal Donkey Serum, Invitrogen, Thermo Scientific). To study proliferation and neurogenesis, serial sections were previously pre-treated in 2M HCl for 15 min at 30°C. Subsequently, brain sections were incubated for 24 h at 4°C in the same pre-incubation stock solution (PB+Triton+Serum) containing different combinations of the following primary antibodies: ab6326 (Abcam) for CldU; AB2253 (Sigma Aldrich) for Doublecortin; ab8898 (Abcam) for Histone H3 (tri methyl K9); ab78517 (Abcam) for Histone H4 (dimethyl K20, tri methyl K20); TYR1336 Antibody (PhosphoSolution) for NMDA GluN2B Subunit (p1516-1336); ab32423 (Abcam) for brain lipid-binding protein (Blbp); 33D3 (Epigentek) for 5-methylcytosine (5mC); 39791(Active-Motif) for 5-Hydromethylcytosine (5hmC) antibody and L1516 (Merck) for Lectin from *Wisteria floribunda* (WFA). After rinsing in PB, the sections were incubated for 2 h at room temperature in the appropriate combinations of Alexa 488-594-647-conjugated anti-mouse/rabbit/guinea pig or rat IgG secondary antibodies (1:500; Molecular Probes, Eugene, OR, USA). Sections were also stained with a nuclear stain DAPI (4,6-diamidino-2-phenylindole; Sigma, St. Louis, MO, USA). In all cases, the sections used were those closest to the injection area and thus in the same positions.

We obtained stitched image stacks from the whole DG at the hippocampus area. These were recorded at 1-μm intervals through separate channels with a 20x lens for the analysis of neurogenesis and the extracellular matrix (ECM) and at 0.45/0.6-μm intervals through separate channels with a 40x lens for the analysis of the N-methyl-D-aspartate (NMDA) GluN2B subunit, and DNA and histone methylation, respectively (Nikon A1R confocal microscope, NA 0.75, refraction index 1, image resolution: 1024 × 1024 pixels). The same range of z-slices was obtained from each slide in each experiment. Adobe Photoshop (CS4) software was used to build the figures.

### 2.7. Immunohistochemical quantifications

Dcx/Blbp/CldU-immunoreactive positive cells were quantified by counting the number of cells localized in each DG, distinguishing between the subgranular zone (SGZ), granular cell layer (GCL), and the adjacent hilus region, in four consecutive medial brain sections. For perineuronal nets (PNN) of the ECM, positive units labeled with WFA were counted in the DG (including the GCL and hilus) from three consecutive medial brain sections. To determine the density of different cells, the DG was traced on the DAPI channel of the z projection of each confocal stack of images, and the area of this structure was measured using the freehand drawing tool in Fiji. This area was multiplied by the stack thickness to calculate the reference volume. The number of positive cells was divided by the reference volume, and the density (number of cells/mm^3^) of cells was calculated.

For the 5mC, 5hmC, H3K9me3, and H4K20me3 antibodies, confocal settings (laser intensity, gain, pinhole) were kept constant for all images, which were captured in the same confocal session. Using the DAPI channel, the whole GCL area was previously selected as a ROI. An invariant subtract background and threshold was then set in Fiji for each stack projection. The area above the threshold was measured in Fiji and the mean fluorescence in the ROI was calculated and compared between vehicle and folate experimental groups.

For the GluN2B and Sox2 antibodies, the quantification was carried out with the ImageJ program by measuring the percentage of area occupied by the receptor immunoreactivity in the molecular layer of the DG and by YF Sox2 immunoreactivity in the hilus region, SGZ, and GCL. An invariant subtract background and threshold were also set in Fiji for each stack projection.

### 2.8. Nuclear fractionation of cultured human cells

The method to fractionate the cell nucleus has been previously reported (Ritter et al., 2018), and the presence of FRα-binding peptide in the nuclear fraction was shown by western blot analysis using the antibody against FRα1-binding peptide (Thermo Fisher, ref: PA5-101588).

### 2.9. Nuclear fractionation of cultured human cells and western blotting

SK-N-SH cells were treated with folate (0.5 or 1mM), FRα-binding peptide (0.5 or 1mM) or Dulbecco’s Modified Eagle Medium (DMEM) for 30 minutes at 37ºC. Total cell lysate or nuclear fraction were obtained as previously reported (Hulin-Curtis et al., 2020). Whole cell lysate or cell nuclear pellets of each experiment were then quantified by the BCA protein assay. Samples were separated on 10% SDS-PAGE and electrophoretically transferred to a nitrocellulose membrane (Schleicher & Schuell GmbH). The membrane was blocked by incubation with 5% semi-fat dried milk in PBS and 0.1% Tween 20 (PBSM), followed by 1-h incubation at room temperature with the primary antibody in PBSM. The following primary antibody dilutions were used: anti-folate receptor alpha 1 (1/300; Thermofisher, ref: PA5-101588) for nuclear fraction and anti-Sox2 (1/1000; R&D systems, ref: AF2018) for total cell lysate. After three washes, the membrane was incubated with a horseradish peroxidase-anti-rabbit Ig conjugate (DAKO), followed by several washes in PBS-Tween 20. The membrane was then incubated for 1 min in Western Lightning reagents (PerkinElmer Life Sciences). Blots were quantified using the EPSON Perfection 1660 scanner and the Image J software.

### 2.10. Behavioral tests

Open field (OF) and novel object recognition (NOR) tests were performed as described previously (Rodriguez-Matellan et al., 2020). Locomotor activity, as well as anxiety and depression-like behavior, was evaluated in old mice using the OF test, which was used as habituation for the NOR test. In brief, on the first day, the mice were placed individually in a 45×45 cm plastic box with vertical opaque walls for 10 min. Each session was recorded, and data like distance, average speed, time immobile, and time on the center square of the box were analyzed with ANY-maze software. On the second day, the mice were placed in the same box for 5 min and allowed to explore two identical objects: A and B (two black rooks, chess piece). The two objects were placed on the long axis of the box each 13 cm from the end of the box. After each exposure, the objects and the box were wiped with 70% ethanol to eliminate odors that could potentially condition the behavior of successive mice. Two hours after the familiarization trial, each mouse was released into the box with the same object previously used (object A) and a new one (object C, a tower of colored plastic pieces), instead of object B (short-term memory test). The position of object C was the same as that of object B in the familiarization trial. The mice were given 5 min to explore the box. Five days later, the mice were released into the box again, with object A in the same position and another new object (object D, a falcon tube filled with wood chips) in the same position as object C in the previous test. Again, the animals were allowed 5 min to explore the box (long-term memory test). They were considered to show recognition when their head was <2 cm from the object, and activity where exploration was less than 1 sec was ruled out.

The memory index (MI) was used to measure recognition memory performance. It was defined as the ratio of time spent exploring or the number of entries in the new object (tC or tD) to the time spent exploring both objects (tA+tC or tA+tD) (MI = [tC/(tA + tC)] X100 and [tD/(tA + tD)] X 100). ANY-maze software was used to calculate the time mice spent exploring the different objects.

To assess spatial memory, a Y maze test based on published protocols with modifications (Kraeuter et al., 2019) was performed. This test is based on the innate curiosity of rodents to explore novel areas. First, the mice were placed into one of the arms of a black Y-maze apparatus, comprising three plastic arms forming a “Y” shape. They were then allowed to explore the maze for 10 min, with one of the arms closed (training trial). After 1 h, the mice were returned to the same arm of the Y maze (start arm) and were allowed to explore all three arms of the maze for 5 min. The number of entries into and the time spent in each arm were registered on video recordings and analyzed by ANY-maze software. We compared the percentage of time spent in the “novel” arm during the whole trial as a measure of spatial memory performance.

### 2.11. Synthetic synthesis of the FR_α_-binding peptide

To mimic the cognitive results produced by intrahippocampal injection of folate but activating only FRα-binding peptide, we used a synthetic FRα-binding peptide (purity of 90%) with the following sequence: CTVRTSAEC (Hulin-Curtis et al., 2020). This peptide was prepared by the company Abyntek (Vizcaya, Spain).

### 2.12. Statistical analysis for immunohistochemical and behavior assessments

The data are presented as mean values ± SD. Statistical analyses were performed using GraphPad Prism 8. To compare the two experimental groups (injected with vehicle and with folate or FRα-binding peptide), an unpaired Student’s t test was carried out. A 95% confidence interval was applied for statistical comparisons.

## 3. Results

### 3.1. Partial improvement in the object recognition and Y maze tests of mice treated with folate but not with S-Adenosyl-Methionine (SAM) or vitamin B12

SAM, folate, or vehicle was intracranially injected (single dose) into the dorsal hippocampal hilar region (**SI 1**) of old mice. The effects of these injections on behavioral tests were analyzed one week later (see Methods). In the first round of surgeries, we screened SAM treatment in a group of 14 mice. SAM injection had no effect on behavioral performance in the Open Field (OF) test (**Fig. 1a**) or memory (**Fig. 1c, e, g**) tests. However, mice treated with folate (n=15; **Fig. 1b, d, f, h**) showed differences in performance in both tests. In this regard, they showed an improvement in locomotor activity and anxiety-depression-like behavior compared to vehicle-treated mice. **Figure 1f** and **1h** shows the differences in long-term memory (5 days) in the novel object recognition and in spatial short-term memory.

**Figure 1.**
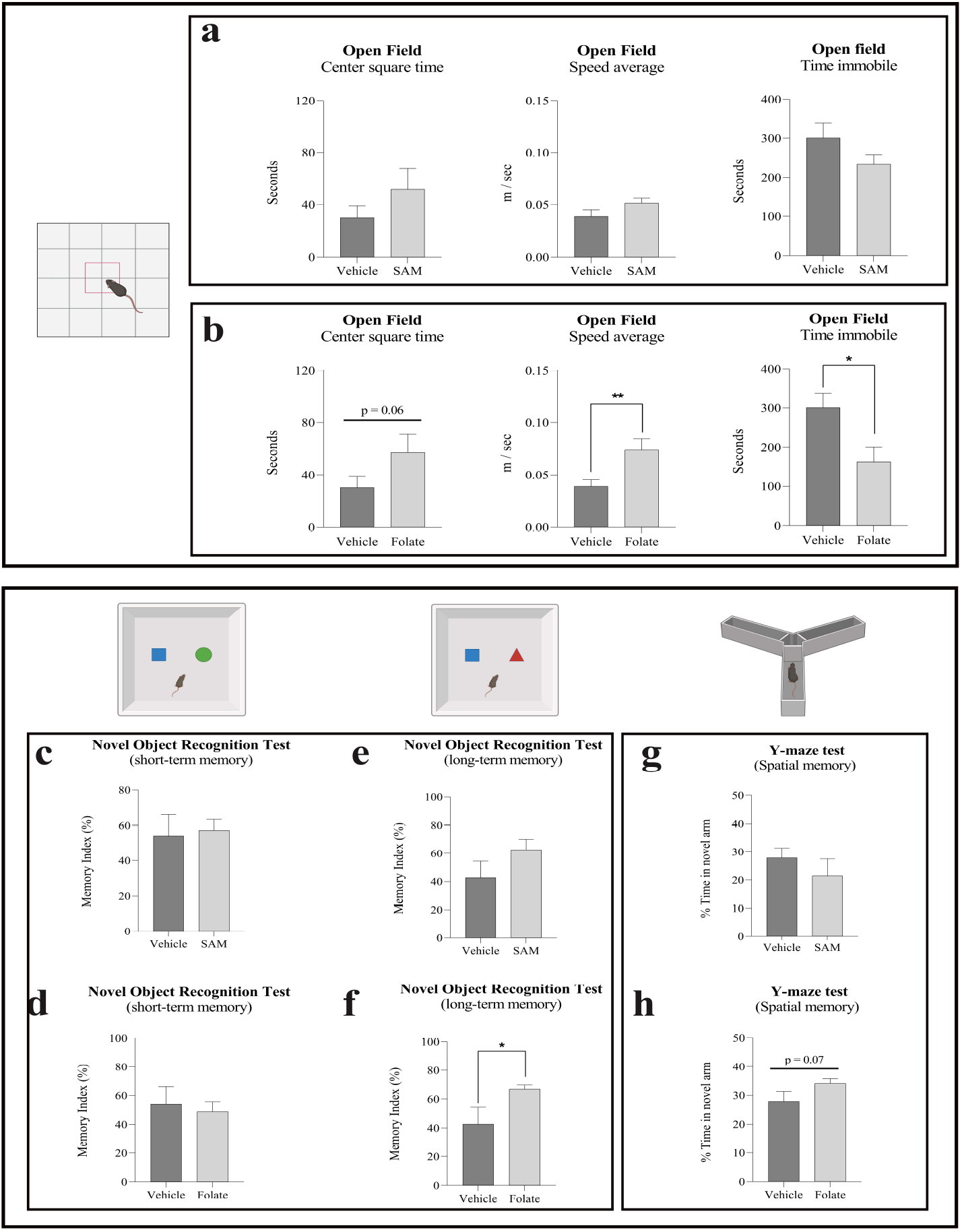
Effect of methyl donors SAM and folate on behavior. Old wild-type mice receiving a single dose of folate via hippocampal infusion showed behavioral changes when compared with mice treated with vehicle solution. This observation was not found for S-adenosylmethionine metabolite (SAM) infusion. (**a, b**) Histograms show some of the main data from the open field test: time spent by mice in the center square of the box, average speed of movement, and time spent immobile. Folate-treated mice showed an improvement in anxiety-depression-like behavior (indicated by the time spent in the center square), as well as in general motor activity (illustrated by the rest of the histograms). (**c-h**) Histograms show results from two memory tests: namely the novel object recognition test (NOR) for short (**c, d**) and long-term recognition memory (**e, f**), and the Y maze test for spatial memory (**g, h**). Significant differences were found in folate-treated mice in comparison with the vehicle-treated group (**f and h**), but none were found between SAM-treated and vehicle-treated mice in any memory test (**c, e, g**). Black bars represent the mean± SD of vehicle-treated groups and gray bars the mean± SD of SAM/Folate-treated groups. All data were analyzed by Student’s t-test. *p<0.05, **p<0.01.

The result of a single injection of folate was similar to that reported for old mice expressing YF in a cyclical manner (Rodriguez-Matellan et al., 2020). Folate improved the long-term memory (5 days) post-familiarization with the novel object (**Fig. 1f**), but no significant differences were found for short-time (2 h) memory (**Fig. 1d**). In addition, folate-treated mice showed improved spatial memory, as reflected by their performance in the Y maze test (**Fig. 1h**). Given our observations that folate injection affected mouse behavior, we examined in greater detail the feasible cellular effects of this compound. To this end, we tested different concentrations of folate. Of note, toxic side effects and even death were observed above 5 mM. We then tested the effect of 2 μl per hemisphere of folate at a concentration of 0.5 mM. This concentration was also used in a previous study (Boshnjaku et al., 2012).

Later, the performance of 12 mice treated with vitamin B12 or vehicle was assessed (**SI 2**). No alteration in locomotor activity (distance traveled, average speed, or time spent immobile) in the OF test or in anxiety-depression-like behavior (time spent in the central square time) was detected in the former group (**SI 2a**). Memory tests (**SI 2b**), focusing on short-term (2 h after training trial) and long-term (5 days after training trial) recognition memory, and on spatial memory did not reveal significant differences between vehicle- and vitamin B12-treated mice.

### 3.2. Possible effect of folate

The effect of folate could occur at different levels related to: a) increase in AHN, b) epigenetic changes in histones or DNA or c) as inducer of transcription factors. In addition, there are several biochemical outputs of folate metabolism (**SI 3**) that will not be studied in this work but could produce some side effects on cells.

In Supplementary Information, the effect of folate on a) adult neurogenesis at the dentate gyrus (**SI 4**) and b) the impact of folate on the methylation of histone and DNA in hippocampal neurons are indicated (**SI 5, 6)**. However, we have focused our studies on c) folate as inducer of transcription factor.

### 3.3. FRα in the cell nucleus could act as a transcription factor

In addition to acting as a methyl donor, folate may exert additional actions on DNA. Upon binding to folate, the folate cell receptor (FRα) (Alam et al., 2019) can be internalized into the cell nucleus and nuclear FRα may act as a transcription factor (Boshnjaku et al., 2012) to facilitate the expression of specific genes (see also **SI 3**).

To test, in a direct way, the action of folate in nuclear internalization of FRα in neuronal cells, we have used a neuronal human cell line (SK-N-SH) as a model to confirm the results reported by Boshnjaku et al. (Boshnjaku et al., 2012) in other cell lines showing that the presence of folate or FRα-binding peptide results in the internalization of FRα into nucleus (**SI 8a**,**b**) of these neuronal cells in a similar fashion to that occurring in other cell types (Boshnjaku et al., 2012). In addition, upon neuronal localization of FRα, we found an increase in the expression of Sox2 protein (**SI 8c, d**), one of the YF. This finding supports a previous report (Mohanty et al., 2016) and could explain how folate could replace YF to facilitate cell rejuvenation.

Also, the expression of Yamanaka factors like Sox2 could increase the expression of GluN2B (Rodriguez-Matellan et al., 2020). These experiments are compatible with a possible effect of folate in the cognitive improvement found in old mice, previously shown in the article.

### 3.4. Effect of folate on transcriptional changes

Furthermore, changes in gene expression occur during aging. Some genes expressed in young cells are no longer expressed, whereas, in old cells, genes that were not previously expressed in young cells could appear. For example, the gene codifying for GluN2B protein is expressed mainly in young neurons (Paoletti et al., 2013), whereas genes involved in the formation of PNN are found mostly in aged tissue (Tanaka and Mizoguchi, 2009). Both of them are structures involved in hippocampal neuroplasticity mechanisms.

#### 3.4.1. Changes in GluN2B expression

We previously reported (Rodriguez-Matellan et al., 2020) that the presence of YF induces an increase in the expression of GluN2B in the molecular layer of the DG. GluN2B is one of the subunits of the N-methyl-D aspartic acid (NMDA) receptor, which is present mainly in young animals and may be related to greater cognitive capacity (Paoletti et al., 2013). **Figure 2** shows the immunoreactivity of GluN2B in the DG (**Fig. 2a, b**) of mice treated or not with folate. GluN2B immunoreactivity in the molecular layer increased in treated animals (**Fig. 2c**)

**Figure 2.**
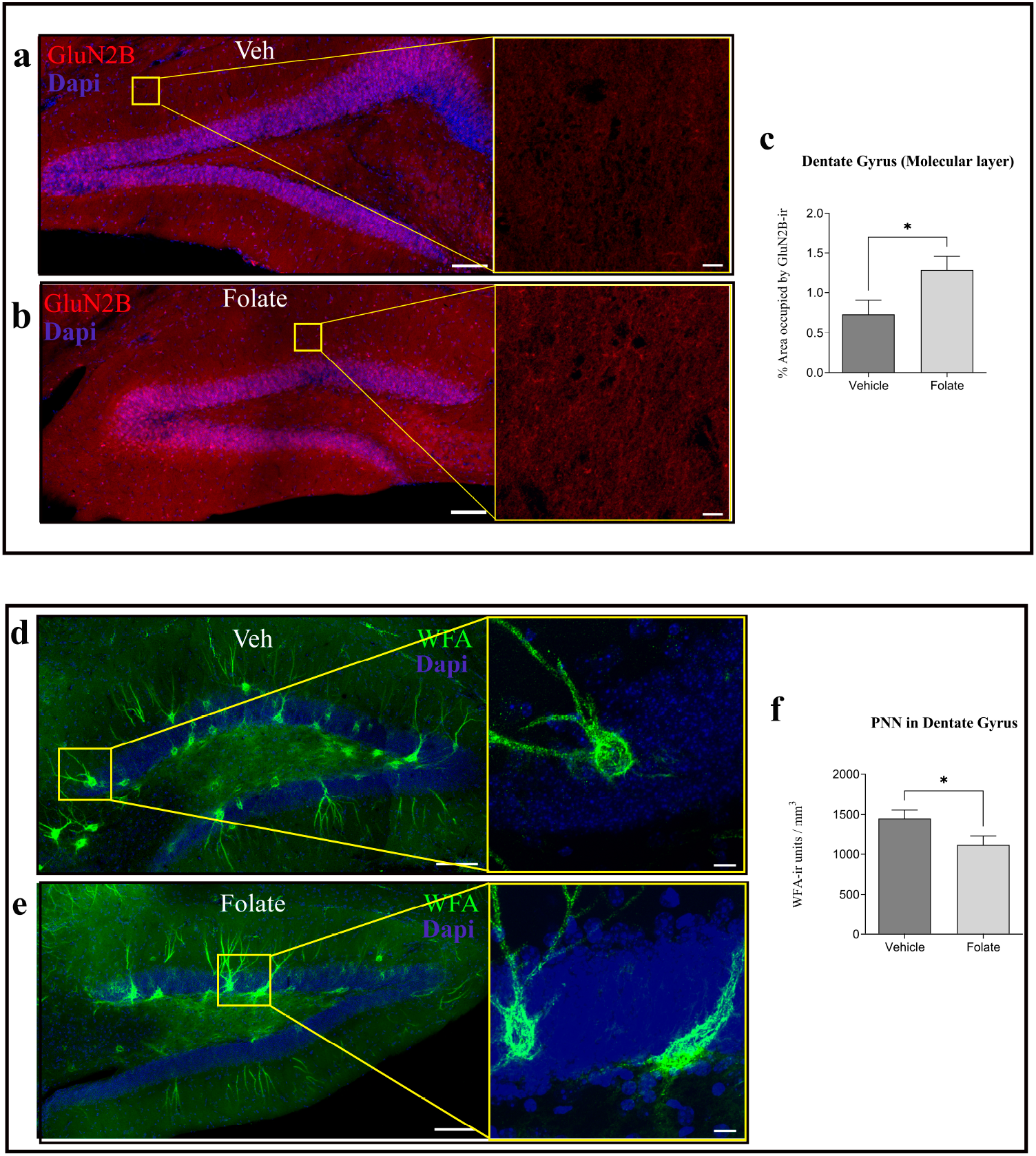
Effect of folate, GluN2B expression and extracellular matrix organization. Effects of folate on the presence of the NMDA receptor subunit GluN2B in mature hippocampal neurons. (**a, b**) Representative stitched confocal images of GluN2B distribution in the dentate gyrus (DG) of old wild-type mice treated with vehicle (**a**) or folate (**b**). Higher magnifications of the molecular cell layer of **a** and **b** (white squares) are shown. (**c**) Histogram showing the percentage of area occupied by GluN2B signal in the molecular layer of the DG. The effect of the absence (black) or presence (gray) of folic acid is shown. Treatment with folate induced a significant increase in GluN2B expression in the DG. **(d, e)** Representative stitched confocal images of Wisteria floribunda agglutinin (WFA, in green), as a marker of perineuronal nets (PNN) in the DG of vehicle-treated and folate-treated mice. Nuclear marker DAPI is shown in blue. Higher magnifications of the molecular cell layer of **d** and **e** (yellow squares) are shown. (**f**) Single-dose treatment with folate resulted in statistically significant changes in PNN density at the DG (Mean ± SD; *p<0.05, Student’s t test). Scale bars, 100 μm (a, b, d, e) and 10 μm (higher magnifications).

#### 3.4.2. Changes in extracellular matrix organization

The ECM is organized into perineuronal nets (PNN). These PNN are involved in synaptic stabilization in the mature brain (Flores and Mendez, 2014). They comprise proteoglycans like neurocan, brevican, aggrecan, phosphacan, and versican, which can be linked to PNN through their binding to hyaluronan (Deepa et al., 2006; Rowlands et al., 2018). These PNN can ensheath interneurons, thereby affecting their function (Wen et al., 2018). Aging is associated with an increase in PNN in some regions of the brain, including the hippocampus (Ueno et al., 2019; Vegh et al., 2014). Thus, we checked the possible reorganization of these structures in response to folate injection in the hippocampal region (**Fig.2d, e**). Folate treatment induced a significant reduction in PNN unit density labeled with Wisteria floribunda agglutinin (WFA) lectin in comparison with vehicle-injected counterparts (**Fig. 2f**).

### 3.5. What is the main effect of folate on cognitive improvement?

To test whether the effect of folate on cognition is due mainly to changes in DNA methylation or to alterations of transcription modulated by factors like nuclear FRα, we tested the effect of an FRα-binding peptide (Hulin-Curtis et al., 2020) to determine whether it can mimic the effect of folate. To this end, we synthesized a peptide (CTVRTSAEC) able to bind FRα (Hulin-Curtis et al., 2020). Although we took into account that the FRα-binding peptide is not a methyl donor and should only facilitate the action of FRα-binding peptide as a transcriptional factor, we first tested, anyway, whether the peptide contributes to DNA methylation. No significant changes in this process were detected (**SI 9**). Thus, we tested whether the peptide induces changes at the transcriptional level. Then, we addressed whether it could also facilitate the internalization of FRα-binding peptide into the cell nucleus of SK-N-SH cultured cells. In this regard, we observed that the peptide facilitated FRα uptake into this cell structure, where it may act as a transcription factor (**SI 8a, b**), increasing the expression of Sox2 in that cell line (**SI 8c, d**).

#### 3.5.1. Effect of the FRα-binding peptide on Sox2 expression

As previously indicated, folate facilitates the expression of YF, including Sox2 (Mohanty et al., 2016). Thus, we studied whether the FRα-binding peptide also affects Sox2 expression in a mouse model. Treatment of aged mice with this peptide resulted in a significant increase in Sox2 expression in various regions of the DG (**Fig. 3a, b, c**).

**Figure 3.**
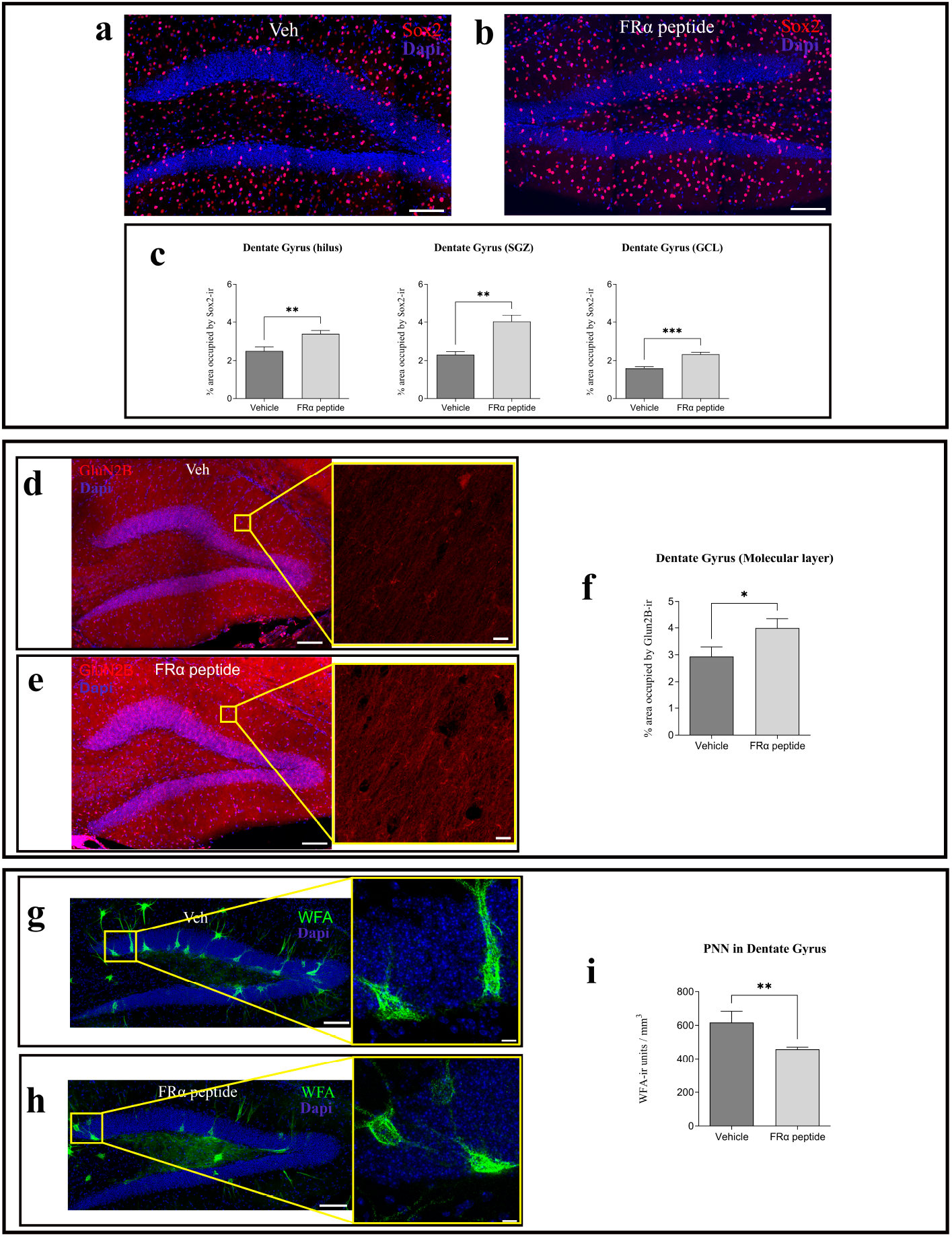
Effect of FRα-binding peptide on the expression of Yamanaka factor Sox2, GluN2B expression and extracellular matrix organization. (**a, b**) Representative stitched confocal images of Sox2 expression (in red) in the dentate gyrus (DG) of vehicle-treated and FRα-treated mice. Nuclear marker DAPI is shown in blue. (**c**) Single-dose treatment with FRα-binding peptide resulted in statistically significant changes in Sox2 immunoreactivity expression in the hilus region, granular cell layer (GCL), and subgranular zone (SGZ) of the DG. It is shown the effects of the FRα-binding peptide on the presence of the NMDA receptor subunit GluN2B in mature hippocampal neurons. (**d, e**) Representative stitched confocal images of GluN2B distribution in the dentate gyrus (DG) of old wild-type mice treated with vehicle (**d**) or FRα-binding peptide (**e**). Higher magnifications of the molecular cell layer of **d** and **e** (yellow squares) are shown. Nuclear marker DAPI is shown in blue. (**f**) Histogram showing the percentage of area occupied by the GluN2B signal in the molecular layer of the DG. The effect of the absence (black) or presence (gray) of FRα is shown. Treatment with FRα-binding peptide induced a significant increase in GluN2B expression in the DG. **(g, h)** Representative stitched confocal images of Wisteria floribunda agglutinin (WFA, in green), as a marker of perineuronal nets (PNN) in the DG of mice injected with vehicle or FRα-binding peptide. Nuclear marker DAPI is shown in blue. Higher magnifications of the molecular cell layer of **g** and **h** (yellow squares) are shown. (**i**) Single-dose treatment with FRα-binding peptide resulted in statistically significant changes in PNN density at the DG. (Mean ± SD; *p<0.05, **p<0.01; Student’s t test). Scale bar, 100 μm (a, c, d, f and g) and 10 μm (higher magnifications).

#### 3.5.2. Effect of the FRα-binding peptide on GLUN2B expression

Given that the presence of YF, such as Sox2, facilitates the expression of GluN2B (Rodriguez-Matellan et al., 2020), we addressed whether the presence of the FRα-binding peptide, which upregulates the expression of Sox2, also increases the expression of GluN2B in an old mouse. Treatment with this peptide led to an increase in GluN2B expression in the molecular layer of the DG (**Fig. 3d, e, f**), as occurred with folate treatment.

#### 3.5.3. Effect of the FRα-binding peptide on extracellular matrix organization

Previously, it was shown that folate treatment caused changes in ECM organization. The FRα-binding peptide affected this organization in a similar fashion to folate, thereby reducing the density of PNN in the DG (**Fig. 3g, h, i**).

Taking into account the results shown in **Figure 3**, we suggest that neuronal signaling is enhanced through changes in excitatory (glutamatergic) (Paoletti et al., 2013; Rodriguez-Matellan et al., 2020) but also in inhibitory neurons (Wen et al., 2018). In addition, both results could be related to each other, since alterations in the expression of PNN components during aging could be modulated by alterations of NMDA function (Yamada et al., 2017).

### 3.6. Cognitive improvement in the presence of FRα-binding peptide

Therefore, we tested whether the treatment with the FRα-binding peptide has a similar effect to that of folate on cognition. Intrahippocampal injection of this peptide under the same conditions as previous folate experiments did not bring about significant changes in short-term memory performance, either in spatial memory or recognition memory, relative to vehicle-treated mice (**Fig. 4a, b**). However, treatment with the FRα-binding peptide led to a significant long-term improvement of recognition memory, thereby reproducing the same results as those achieved with folate injections (**Fig. 4c**).

**Figure 4.**
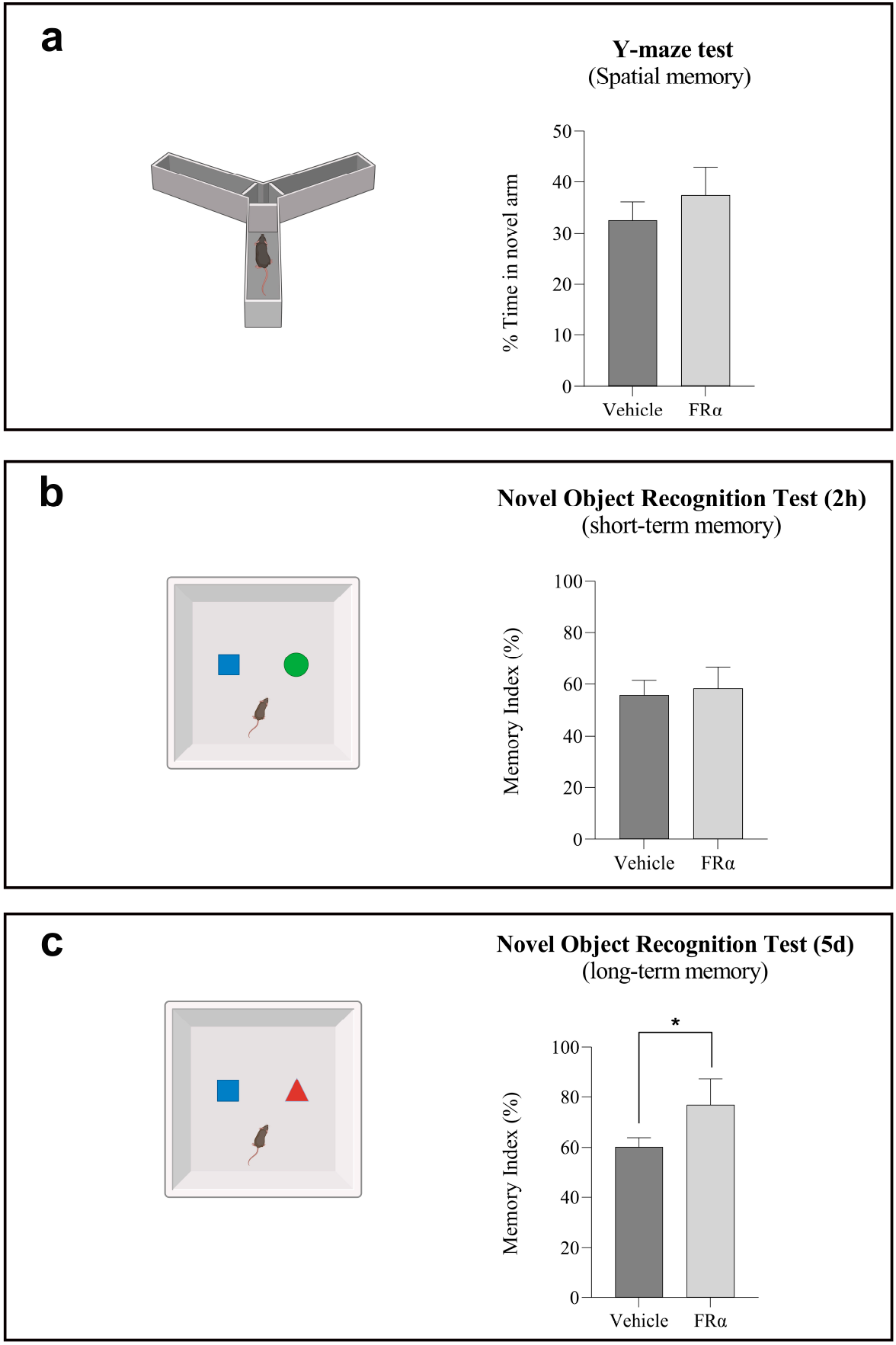
Effect of FRα-binding peptide on cognition. Histograms show results from two memory tests: the Y-maze test for spatial memory (**a**) and the novel object recognition (NOR) for short-(2 hours) (**b**) and long-term recognition memory (5 days) (**c**). Mice treated with the peptide showed a significant improvement in memory retention after 5 days since first training, as shown by memory index of novel object recognition test. Treatment with FRα-binding peptide did not have a significant effect on short-term recognition memory. Black bars represent mean ± SD of the vehicle-treated group and gray bars represent mean± SD of the peptide-treated group. All data were analyzed by Student’s t-test. *p<0.05

## 4. Discussion

Here we devised and tested a novel strategy, involving a simple compound, to replace the rejuvenating effects of YF on old cells that make them acquire the traits of younger ones (partial reprogramming). We found that a single injection of folate was a suitable alternative to cyclic treatment with YF. First, folate treatment resulted in epigenetic changes related to the rejuvenation of methylation patterns, as achieved with YF. Also, we observed an increase in the expression of subunit GluN2B of the NMDA receptor (Rodriguez-Matellan et al., 2020; Zhang et al., 2015). This subunit is present mainly in excitatory neurons of young animals (Tang et al., 1999) and this increase may imply enhanced synaptic plasticity, as previously described for YF expression (Rodriguez-Matellan et al., 2020). The increased plasticity induced by the transient presence of folate could underlie the improvement in recognition and spatial memory observed in these old mice. Interestingly, long-term but not short-term recognition memory was enhanced by folate treatment, as occurred with YF cyclical overexpression (Rodriguez-Matellan et al., 2020). Furthermore, aging and age-associated diseases like Alzheimer disease have been related to an increase in perineuronal nets (PNN) in different regions of the brain, including the hippocampus (Karetko-Sysa et al., 2014; Tanaka and Mizoguchi, 2009; Ueno et al., 2019; Vegh et al., 2014; Yamada et al., 2017). PNN are a specific type of ECM wrapped around some interneurons (Wen et al., 2018) that restrict the plasticity of these cells. PNN are dynamic structures that undergo changes throughout postnatal development, and their reduction has been associated with the facilitation of neuronal plasticity and memory enhancement (Romberg et al., 2013; Rowlands et al., 2018; Thompson et al., 2018). In this regard, our results in aged mice showing the reorganization of the ECM by reducing the density of PNN replicate those reported for the rejuvenating effects of YF on the hippocampus (Rodriguez-Matellan et al., 2020).

The mechanism for YF partial reprogramming could involve transcriptional changes and an increase in DNA methylation (Lopez-Bertoni et al., 2015; Tsai et al., 2012; Wu et al., 2018). Given that the latter process can also be enhanced by the presence of methyl donors or cofactors, we tested the possible partial reprogramming effect of folate and SAM as methyl donor and of vitamin B12 as a cofactor. Although methionine is the universal methyl donor, folate may act as a donor through folate-methionine metabolism (Fenech, 2010). However, some differences between the presence of methionine and folate can be found. Toxicity for high levels of methionine but not for folate were found in humans (Garlick, 2006). Methionine toxicity could be due to the fact that methionine is a precursor of the toxic homocysteine (Perna et al., 2003). Also, it should be noted that a high-methionine diet induces Alzheimer disease-like symptoms (Pi et al., 2021). On the other hand, other side effects can occur; for example, if methionine is converted to SAM, the latter can also re-methylate methionine through homocysteine (Fenech, 2010; Ostrakhovitch and Tabibzadeh, 2019). Also, there is a decrease in the levels of folate and an increase in homocysteine with age (Fenech, 2010). In addition, low levels of folate may correlate with the cognitive decline of elderly patients (Araujo et al., 2015). In fact, folate supplement has been proposed as a potential treatment for improving memory deficits (Cummings et al., 2017; Fioravanti et al., 1998). On the other hand, the link between vitamin B12, folate, and homocysteine and cognitive performance during aging is well known (Riggs et al., 1996).

Of note, despite the involvement of both folate and SAM as possible methyl donors, only folate was found to have a similar impact on cognition as YF. This difference could be explained by the interaction of this compound with the folate receptors, FRα being the most relevant. FRα is a transcription factor that is involved in the expression of the YF Sox2 (Mohanty et al., 2016). The same long-term recognition memory enhancement induced by treatment with FRα-binding peptide as with folate injections would also support this hypothesis.

Folate deficiency related to aging may result in the exposure of hypomethylated DNA present in heterochromatin regions, after despiralization of chromatin (Fenech, 2010). In this way, aging has been related to a loss of DNA methylation (Bollati et al., 2009; Wilson and Jones, 1983), particularly at constitutive heterochromatin (Tsurumi and Li, 2012; Villeponteau, 1997; Wang et al., 2016), which may potentially lead to altered chromatin functioning and genomic instability (Hu et al., 2014). In this regard, in the model of aging due to the loss of heterochromatin, heterochromatin markers like DNA methylation decrease with age at the time of heterochromatin loss (Villeponteau, 1997). Otherwise, aging could be also related to an increase in DNA methylation, mainly in euchromatin regions (Horvath and Raj, 2018). Indeed, loss of heterochromatin has been found in premature aging disorders like Hutchinson-Gilford progeria (Scaffidi and Misteli, 2005; Shumaker et al., 2006) and Werner syndrome (Zhang et al., 2015). Heterochromatin markers like DNA cytosine-5 methylation are globally decreased in the hippocampus during aging (Chen et al., 2012; Szulwach et al., 2011), leading to general DNA hypomethylation (Ciccarone et al., 2018). The latter may facilitate the presence of aging signatures (Johansson et al., 2013; Zampieri et al., 2015), but if DNA methylation is promoted, these signatures may be corrected. However, the main action of folate could be the regulation of the expression of specific genes. Although folate and Frα peptide share that effect on gene expression, folate, together with its nine derivatives (Nzila et al., 2005), but not Frα peptide, is involved in other many varied biological actions (**SI 3**). For example, supplying one carbon units (methyl groups) for several metabolic pathways like the biosynthesis of methionine, the biosynthesis of purines and thymidine (related to DNA synthesis), aminoacid homeostasis (Annibal et al., 2021; Bailey and Gregory, 1999), epigenetic maintenance (as methyl donor) or for redox defense (Ho et al., 2015). In this work, we have indicated that likely the main effect of folate on cognition is not anyone of the previous effects as methyl donor but it is due to its action on FRα, shared with FRα binding peptides, which can increase the level of factors like Sox2 and GluN2B or produce changes in the organization of the ECM, specifically PNN, which, in addition, will improve cognitive functions such as long-term memory. Thus, our current working hypothesis indicates a possible pathway to explain the role of folate or the FRα-binding peptide in cognitive improvement of old mice, described in the following pathway: **folate or peptide, nuclear FR**α, **Sox2**↑, **GluN2B** ↑, **ECM organization** ↓, **improved cognition**.

We observed several similarities and differences between the addition of folate and FR-binding peptide in our working hypothesis (**Fig. 5**). In this context, we propose that folate and/or the indicated FRα-binding peptide promote cognition, mainly through the action of nuclear FRα. However, for folate, we cannot rule out that changes at the DNA methylation level may help to facilitate this functional improvement. In fact, folate treatment led not only to enhanced recognition memory but also better spatial memory when compared to the results of the mice treated with the FRα-binding peptide. On the other hand, we have not found, up to now, any negative side effects caused by treatment with this peptide. Based on the previous comments, to avoid possible side effects for potential therapies could be more suitable the use of FRα binding peptide than that of folate.

**Figure 5.**
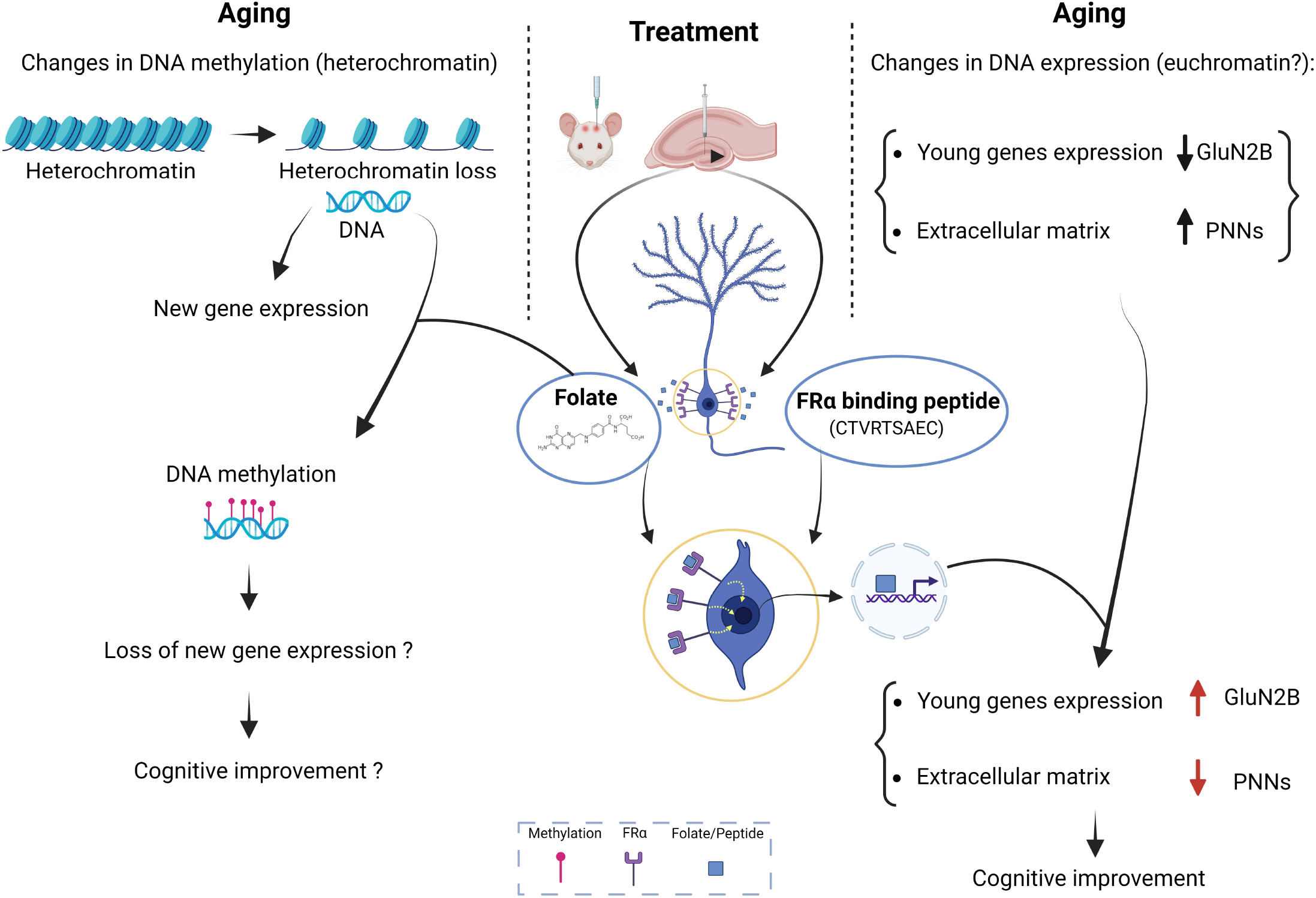
Scheme of main results. Left-According to the heterochromatin loss aging model (Villeponteau, 1997), heterochromatin biomarkers like DNA methylation (in heterochromatin regions) are lost with age. Right-At the same time, during aging, the brain undergoes changes in DNA expression, leading to decreased levels of certain genes, like GluN2B, related with more juvenile states or with increased levels of other structures like perineuronal nets (PNN). Center-In this work, we report that folate and the FRα-binding peptide may facilitate the transport of FRα-binding peptide to the cell nucleus, where the receptor acts like a transcription factor (Boshnjaku et al., 2012). Consequently, treatment with either of these two compounds (folate or FRα-binding peptide) reverted the age-associated loss of neuroplasticity by changing expression in GluN2B and in PNN units, thereby leading to cognitive improvement. Furthermore, treatment with folate but not with the FRα-binding peptide led to a significant increase in DNA methylation, thereby recovering the heterochromatin loss associated with aging.

In summary, of the three possible effects of folate on cognitive improvement tested: a) increasing trend in adult neurogenesis, b) increase in DNA methylation or c) increase in neuroplasticity by reorganizing extracellular matrix structures or rising the expression of juvenile genes like GluN2B, we suggest that effect c) could play the main role in cognitive improvement.

## Conclusions

In conclusion, our results indicate that folate or FRα-peptide binding treatment can induce rejuvenation of aged DG cells enhancing cognitive improvement. Thus, FRα peptide or folate through the activation of FRα pathway, emerge as an alternative strategy in the central nervous system to treat cerebral aging and age-associated neurodegenerative diseases.

## List of abbreviations

AHN: Adult hippocampal neurogenesis
DG: Dentate gyrus
DMEM: Dulbecco’s Modified Eagle Medium
DNMT: DNA methyltransferase
ECM: Extracellular matrix
FRα: Folate cell receptor
GCL: Granular cell layer
iPSCs: Induced pluripotent stem cells
MI: Memory index
NOR: Novel object recognition
NMDA: N-methyl-D-aspartate
OF: Open field
PBS: Phosphate buffered saline
PNN: Perineuronal nets
SAM: S-adenosyl-methionine
SGZ: Subgranular zone
WFA: Wisteria floribunda agglutinin
YF: Yamanaka Factors

## Data Availability Statement

The data that support the findings of this study are openly available in figshare.com at https://figshare.com/articles/dataset/All_dataset/22277158.

## Author contributions

JA, AAF, and FH designed the experiment. AAF performed animal surgeries, intracerebral injections and behavioral test. RPC carried out support tasks for the maintenance and improvement of animal welfare. RC performed cell cultures and western blots. AAF, FH and JA analyzed the data and drafted the manuscript. AAF, RC, RPC, FH and JA read and approved the final manuscript.

## Acknowledgements

This work has been supported by grants from the Spanish Ministry of Economy and competitiveness: PGC-2018-09177-B-100 (J.A.), PID2020-113204GB-I00 (F.H.) and PID2021-123859OB-100 from MCIN /AEI/10.13039/501100011033 / FEDER, UE (J.A.). This work was supported by CSIC through an intramural grant (201920E104) (J.A.). The Centro de Biología Molecular Severo Ochoa (CBMSO) is a *Severo Ochoa* Center of Excellence (MICIN, award CEX2021-001154-S).

We would like to acknowledge the comments, corrections and advice given by Prof. Carlos López-Otín. We thank Ms. Nuria de la Torre Alonso for technical and editorial assistance. We also thank the Optical and Confocal Microscopy Facility (SMOC) at CBMSO-CSIC for technical support. Graphical abstract, scheme Figure 5 and supplementary Figure 3 were created with BioRender (BioRender.com).

## Ethic declarations

All applicable international, national, and institutional animal welfare guidelines were followed for the experiments involving experimentation animals. All procedures were in accordance with ethical standards and were approved by the pertinent Ethics Committee. Mice were bred in the animal facility of Centro de Biología Molecular Severo Ochoa. They were housed in a specific pathogen-free colony facility under standard laboratory conditions, following European Community Guidelines (directive 86/609/EEC), and handled in accordance with European and local animal care protocols (PROEX 102.0/21). 4–5 mice were housed per cage with food and water available ad libitum, and maintained in a temperature-controlled environment on a 12h/12h light/dark cycle with light onset at 8 a.m.

## Reference list

Alam, C., Aufreiter, S., Georgiou, C.J., Hoque, M.T., Finnell, R.H., O’Connor, D.L., Goldman, I.D., Bendayan, R., 2019. Upregulation of reduced folate carrier by vitamin D enhances brain folate uptake in mice lacking folate receptor alpha. Proc Natl Acad Sci U S A 116(35), 17531–17540.

Annibal, A., Tharyan, R.G., Schonewolff, M.F., Tam, H., Latza, C., Auler, M.M.K., Gronke, S., Partridge, L., Antebi, A., 2021. Regulation of the one carbon folate cycle as a shared metabolic signature of longevity. Nat Commun 12(1), 3486.

Araujo, J.R., Martel, F., Borges, N., Araujo, J.M., Keating, E., 2015. Folates and aging: Role in mild cognitive impairment, dementia and depression. Ageing Res Rev 22, 9–19.

Bailey, L.B., Gregory, J.F., 3rd, 1999. Folate metabolism and requirements. J Nutr 129(4), 779–782.

Benayoun, B.A., Pollina, E.A., Brunet, A., 2015. Epigenetic regulation of ageing: linking environmental inputs to genomic stability. Nat Rev Mol Cell Biol 16(10), 593–610.

Bollati, V., Schwartz, J., Wright, R., Litonjua, A., Tarantini, L., Suh, H., Sparrow, D., Vokonas, P., Baccarelli, A., 2009. Decline in genomic DNA methylation through aging in a cohort of elderly subjects. Mech Ageing Dev 130(4), 234–239.

Boshnjaku, V., Shim, K.W., Tsurubuchi, T., Ichi, S., Szany, E.V., Xi, G., Mania-Farnell, B., McLone, D.G., Tomita, T., Mayanil, C.S., 2012. Nuclear localization of folate receptor alpha: a new role as a transcription factor. Sci Rep 2, 980.

Chen, H., Dzitoyeva, S., Manev, H., 2012. Effect of aging on 5-hydroxymethylcytosine in the mouse hippocampus. Restor Neurol Neurosci 30(3), 237–245.

Chen, R., Skutella, T., 2022. Synergistic Anti-Ageing through Senescent Cells Specific Reprogramming. Cells 11(5), 830.

Chondronasiou, D., Gill, D., Mosteiro, L., Urdinguio, R.G., Berenguer-Llergo, A., Aguilera, M., Durand, S., Aprahamian, F., Nirmalathasan, N., Abad, M., Martin-Herranz, D.E., Stephan-Otto Attolini, C., Prats, N., Kroemer, G., Fraga, M.F., Reik, W., Serrano, M., 2022. Multi-omic rejuvenation of naturally aged tissues by a single cycle of transient reprogramming. Aging Cell 21(3), e13578.

Ciccarone, F., Tagliatesta, S., Caiafa, P., Zampieri, M., 2018. DNA methylation dynamics in aging: how far are we from understanding the mechanisms? Mech Ageing Dev 174, 3–17.

Cummings, J., Scheltens, P., McKeith, I., Blesa, R., Harrison, J.E., Bertolucci, P.H., Rockwood, K., Wilkinson, D., Wijker, W., Bennett, D.A., Shah, R.C., 2017. Effect Size Analyses of Souvenaid in Patients with Alzheimer’s Disease. J Alzheimers Dis 55(3), 1131–1139.

Deepa, S.S., Carulli, D., Galtrey, C., Rhodes, K., Fukuda, J., Mikami, T., Sugahara, K., Fawcett, J.W., 2006. Composition of perineuronal net extracellular matrix in rat brain: a different disaccharide composition for the net-associated proteoglycans. J Biol Chem 281(26), 17789–17800.

Ducker, G.S., Rabinowitz, J.D., 2017. One-Carbon Metabolism in Health and Disease. Cell Metab 25(1), 27–42.

Fenech, M., 2010. Folate, DNA damage and the aging brain. Mech Ageing Dev 131(4), 236–241.

Fioravanti, M., Ferrario, E., Massaia, M., Cappa, G., Rivolta, G., Grossi, E., Buckley, A.E., 1998. Low folate levels in the cognitive decline of elderly patients and the efficacy of folate as a treatment for improving memory deficits. Arch Gerontol Geriatr 26(1), 1–13.

Flores, C.E., Mendez, P., 2014. Shaping inhibition: activity dependent structural plasticity of GABAergic synapses. Front Cell Neurosci 8, 327.

Garlick, P.J., 2006. Toxicity of methionine in humans. J Nutr 136(6 Suppl), 1722S–1725S.

Ho, C.T., Shang, H.S., Chang, J.B., Liu, J.J., Liu, T.Z., 2015. Folate deficiency-triggered redox pathways confer drug resistance in hepatocellular carcinoma. Oncotarget 6(28), 26104–26118.

Horvath, S., Raj, K., 2018. DNA methylation-based biomarkers and the epigenetic clock theory of ageing. Nat Rev Genet 19(6), 371–384.

Hou, P., Li, Y., Zhang, X., Liu, C., Guan, J., Li, H., Zhao, T., Ye, J., Yang, W., Liu, K., Ge, J., Xu, J., Zhang, Q., Zhao, Y., Deng, H., 2013. Pluripotent stem cells induced from mouse somatic cells by small-molecule compounds. Science 341(6146), 651–654.

Hu, Z., Chen, K., Xia, Z., Chavez, M., Pal, S., Seol, J.H., Chen, C.C., Li, W., Tyler, J.K., 2014. Nucleosome loss leads to global transcriptional up-regulation and genomic instability during yeast aging. Genes Dev 28(4), 396–408.

Hulin-Curtis, S.L., Davies, J.A., Nestic, D., Bates, E.A., Baker, A.T., Cunliffe, T.G., Majhen, D., Chester, J.D., Parker, A.L., 2020. Identification of folate receptor alpha (FRalpha) binding oligopeptides and their evaluation for targeted virotherapy applications. Cancer Gene Ther 27(10-11), 785–798.

Johansson, A., Enroth, S., Gyllensten, U., 2013. Continuous Aging of the Human DNA Methylome Throughout the Human Lifespan. PLoS One 8(6), e67378.

Karetko-Sysa, M., Skangiel-Kramska, J., Nowicka, D., 2014. Aging somatosensory cortex displays increased density of WFA-binding perineuronal nets associated with GAD-negative neurons. Neuroscience 277, 734–746.

Kim, J., Lengner, C.J., Kirak, O., Hanna, J., Cassady, J.P., Lodato, M.A., Wu, S., Faddah, D.A., Steine, E.J., Gao, Q., Fu, D., Dawlaty, M., Jaenisch, R., 2011. Reprogramming of postnatal neurons into induced pluripotent stem cells by defined factors. Stem Cells 29(6), 992–1000.

Kraeuter, A.K., Guest, P.C., Sarnyai, Z., 2019. The Y-Maze for Assessment of Spatial Working and Reference Memory in Mice. Methods Mol Biol 1916, 105–111.

Lopez-Bertoni, H., Lal, B., Li, A., Caplan, M., Guerrero-Cazares, H., Eberhart, C.G., Quinones-Hinojosa, A., Glas, M., Scheffler, B., Laterra, J., Li, Y., 2015. DNMT-dependent suppression of microRNA regulates the induction of GBM tumor-propagating phenotype by Oct4 and Sox2. Oncogene 34(30), 3994–4004.

Lopez-Otin, C., Blasco, M.A., Partridge, L., Serrano, M., Kroemer, G., 2013. The hallmarks of aging. Cell 153(6), 1194–1217.

Mohanty, V., Shah, A., Allender, E., Siddiqui, M.R., Monick, S., Ichi, S., Mania-Farnell, B. D G.M., Tomita, T., Mayanil, C.S., 2016. Folate Receptor Alpha Upregulates Oct4, Sox2 and Klf4 and Downregulates miR-138 and miR-let-7 in Cranial Neural Crest Cells. Stem Cells 34(11), 2721–2732.

Nzila, A., Ward, S.A., Marsh, K., Sims, P.F., Hyde, J.E., 2005. Comparative folate metabolism in humans and malaria parasites (part II): activities as yet untargeted or specific to Plasmodium. Trends Parasitol 21(7), 334–339.

Ocampo, A., Reddy, P., Martinez-Redondo, P., Platero-Luengo, A., Hatanaka, F., Hishida, T., Li, M., Lam, D., Kurita, M., Beyret, E., Araoka, T., Vazquez-Ferrer, E., Donoso, D., Roman, J.L., Xu, J., Rodriguez Esteban, C., Nunez, G., Nunez Delicado, E., Campistol, J.M., Guillen, I., Guillen, P., Izpisua Belmonte, J.C., 2016. In Vivo Amelioration of Age-Associated Hallmarks by Partial Reprogramming. Cell 167(7), 1719–1733 e1712.

Olova, N., Simpson, D.J., Marioni, R.E., Chandra, T., 2019. Partial reprogramming induces a steady decline in epigenetic age before loss of somatic identity. Aging Cell 18(1), e12877.

Ostrakhovitch, E.A., Tabibzadeh, S., 2019. Homocysteine and age-associated disorders. Ageing Res Rev 49, 144–164.

Paoletti, X., Oba, K., Bang, Y.J., Bleiberg, H., Boku, N., Bouche, O., Catalano, P., Fuse, N., Michiels, S., Moehler, M., Morita, S., Ohashi, Y., Ohtsu, A., Roth, A., Rougier, P., Sakamoto, J., Sargent, D., Sasako, M., Shitara, K., Thuss-Patience, P., Van Cutsem, E., Burzykowski, T., Buyse, M., group, G., 2013. Progression-free survival as a surrogate for overall survival in advanced/recurrent gastric cancer trials: a meta-analysis. J Natl Cancer Inst 105(21), 1667–1670.

Perna, A.F., Ingrosso, D., Lombardi, C., Acanfora, F., Satta, E., Cesare, C.M., Violetti, E., Romano, M.M., De Santo, N.G., 2003. Possible mechanisms of homocysteine toxicity. Kidney Int Suppl(84), S137–140.

Pi, T., Wei, S., Jiang, Y., Shi, J.S., 2021. High Methionine Diet-Induced Alzheimer’s Disease like Symptoms Are Accompanied by 5-Methylcytosine Elevated Levels in the Brain. Behav Neurol 2021, 6683318.

Planello, A.C., Ji, J., Sharma, V., Singhania, R., Mbabaali, F., Muller, F., Alfaro, J.A., Bock, C., De Carvalho, D.D., Batada, N.N., 2014. Aberrant DNA methylation reprogramming during induced pluripotent stem cell generation is dependent on the choice of reprogramming factors. Cell Regen 3(1), 4.

Pollina, E.A., Brunet, A., 2011. Epigenetic regulation of aging stem cells. Oncogene 30(28), 3105–3126.

Riggs, K.M., Spiro, A., 3rd, Tucker, K., Rush, D., 1996. Relations of vitamin B-12, vitamin B-6, folate, and homocysteine to cognitive performance in the Normative Aging Study. Am J Clin Nutr 63(3), 306–314.

Ritter, M.L., Avila, J., Garcia-Escudero, V., Hernandez, F., Perez, M., 2018. Frontotemporal Dementia-Associated N279K Tau Mutation Localizes at the Nuclear Compartment. Front Cell Neurosci 12, 202.

Rodriguez-Matellan, A., Alcazar, N., Hernandez, F., Serrano, M., Avila, J., 2020. In Vivo Reprogramming Ameliorates Aging Features in Dentate Gyrus Cells and Improves Memory in Mice. Stem Cell Reports 15(5), 1056–1066.

Romberg, C., Yang, S., Melani, R., Andrews, M.R., Horner, A.E., Spillantini, M.G., Bussey, T.J., Fawcett, J.W., Pizzorusso, T., Saksida, L.M., 2013. Depletion of perineuronal nets enhances recognition memory and long-term depression in the perirhinal cortex. J Neurosci 33(16), 7057–7065.

Rowlands, D., Lensjo, K.K., Dinh, T., Yang, S., Andrews, M.R., Hafting, T., Fyhn, M., Fawcett, J.W., Dick, G., 2018. Aggrecan Directs Extracellular Matrix-Mediated Neuronal Plasticity. J Neurosci 38(47), 10102–10113.

Sarkar, T.J., Quarta, M., Mukherjee, S., Colville, A., Paine, P., Doan, L., Tran, C.M., Chu, C.R., Horvath, S., Qi, L.S., Bhutani, N., Rando, T.A., Sebastiano, V., 2020. Transient non-integrative expression of nuclear reprogramming factors promotes multifaceted amelioration of aging in human cells. Nat Commun 11(1), 1545.

Scaffidi, P., Misteli, T., 2005. Reversal of the cellular phenotype in the premature aging disease Hutchinson-Gilford progeria syndrome. Nat Med 11(4), 440–445.

Sen, P., Shah, P.P., Nativio, R., Berger, S.L., 2016. Epigenetic Mechanisms of Longevity and Aging. Cell 166(4), 822–839.

Shumaker, D.K., Dechat, T., Kohlmaier, A., Adam, S.A., Bozovsky, M.R., Erdos, M.R., Eriksson, M., Goldman, A.E., Khuon, S., Collins, F.S., Jenuwein, T., Goldman, R.D., 2006. Mutant nuclear lamin A leads to progressive alterations of epigenetic control in premature aging. Proc Natl Acad Sci U S A 103(23), 8703–8708.

Szulwach, K.E., Li, X., Li, Y., Song, C.X., Wu, H., Dai, Q., Irier, H., Upadhyay, A.K., Gearing, M., Levey, A.I., Vasanthakumar, A., Godley, L.A., Chang, Q., Cheng, X., He, C., Jin, P., 2011. 5-hmC-mediated epigenetic dynamics during postnatal neurodevelopment and aging. Nat Neurosci 14(12), 1607–1616.

Takahashi, K., Yamanaka, S., 2006. Induction of pluripotent stem cells from mouse embryonic and adult fibroblast cultures by defined factors. Cell 126(4), 663–676.

Tanaka, Y., Mizoguchi, K., 2009. Influence of aging on chondroitin sulfate proteoglycan expression and neural stem/progenitor cells in rat brain and improving effects of a herbal medicine, yokukansan. Neuroscience 164(3), 1224–1234.

Tang, Y.P., Shimizu, E., Dube, G.R., Rampon, C., Kerchner, G.A., Zhuo, M., Liu, G., Tsien, J.Z., 1999. Genetic enhancement of learning and memory in mice. Nature 401(6748), 63–69.

Thompson, E.H., Lensjo, K.K., Wigestrand, M.B., Malthe-Sorenssen, A., Hafting, T., Fyhn, M., 2018. Removal of perineuronal nets disrupts recall of a remote fear memory. Proc Natl Acad Sci U S A 115(3), 607–612.

Tsai, C.C., Su, P.F., Huang, Y.F., Yew, T.L., Hung, S.C., 2012. Oct4 and Nanog directly regulate Dnmt1 to maintain self-renewal and undifferentiated state in mesenchymal stem cells. Mol Cell 47(2), 169–182.

Tsurumi, A., Li, W.X., 2012. Global heterochromatin loss: a unifying theory of aging? Epigenetics 7(7), 680–688.

Ueno, H., Suemitsu, S., Murakami, S., Kitamura, N., Wani, K., Takahashi, Y., Matsumoto, Y., Okamoto, M., Ishihara, T., 2019. Alteration of Extracellular Matrix Molecules and Perineuronal Nets in the Hippocampus of Pentylenetetrazol-Kindled Mice. Neural Plast 2019, 8924634.

Vegh, M.J., Rausell, A., Loos, M., Heldring, C.M., Jurkowski, W., van Nierop, P., Paliukhovich, I., Li, K.W., del Sol, A., Smit, A.B., Spijker, S., van Kesteren, R.E., 2014. Hippocampal extracellular matrix levels and stochasticity in synaptic protein expression increase with age and are associated with age-dependent cognitive decline. Mol Cell Proteomics 13(11), 2975–2985.

Villeponteau, B., 1997. The heterochromatin loss model of aging. Exp Gerontol 32(4-5), 383–394.

Wang, J., Jia, S.T., Jia, S., 2016. New Insights into the Regulation of Heterochromatin. Trends Genet 32(5), 284–294.

Watanabe, A., Yamada, Y., Yamanaka, S., 2013. Epigenetic regulation in pluripotent stem cells: a key to breaking the epigenetic barrier. Philos Trans R Soc Lond B Biol Sci 368(1609), 20120292.

Wen, T.H., Binder, D.K., Ethell, I.M., Razak, K.A., 2018. The Perineuronal ‘Safety’ Net? Perineuronal Net Abnormalities in Neurological Disorders. Front Mol Neurosci 11, 270.

Wilson, V.L., Jones, P.A., 1983. DNA methylation decreases in aging but not in immortal cells. Science 220(4601), 1055–1057.

Wu, F., Wu, Q., Li, D., Zhang, Y., Wang, R., Liu, Y., Li, W., 2018. Oct4 regulates DNA methyltransferase 1 transcription by direct binding of the regulatory element. Cell Mol Biol Lett 23, 39.

Yamada, J., Ohgomori, T., Jinno, S., 2017. Alterations in expression of Cat-315 epitope of perineuronal nets during normal ageing, and its modulation by an open-channel NMDA receptor blocker, memantine. J Comp Neurol 525(9), 2035–2049.

Yamanaka, S., 2008. Induction of pluripotent stem cells from mouse fibroblasts by four transcription factors. Cell Prolif 41 Suppl 1(Suppl 1), 51–56.

Zampieri, M., Ciccarone, F., Calabrese, R., Franceschi, C., Burkle, A., Caiafa, P., 2015. Reconfiguration of DNA methylation in aging. Mech Ageing Dev 151, 60–70.

Zhang, W., Li, J., Suzuki, K., Qu, J., Wang, P., Zhou, J., Liu, X., Ren, R., Xu, X., Ocampo, A., Yuan, T., Yang, J., Li, Y., Shi, L., Guan, D., Pan, H., Duan, S., Ding, Z., Li, M., Yi, F., Bai, R., Wang, Y., Chen, C., Yang, F., Li, X., Wang, Z., Aizawa, E., Goebl, A., Soligalla, R.D., Reddy, P., Esteban, C.R., Tang, F., Liu, G.H., Belmonte, J.C., 2015. Aging stem cells. A Werner syndrome stem cell model unveils heterochromatin alterations as a driver of human aging. Science 348(6239), 1160–1163.

