## Supplementary Information for "The role of folate receptor α in the partial rejuvenation of dentate gyrus cells. Improvement of cognitive function in elderly mice"

Antón-Fernández A[1]; Cuadros R[1,2]; Peinado-Cahuchola R[1]; Hernández F[1]; Avila J[1,2*]
[1] Centro de Biología Molecular "Severo Ochoa", CSIC/UAM, Universidad Autónoma de Madrid, Cantoblanco, 28049 Madrid, Spain.
[2] Center for Networked Biomedical Research on Neurodegenerative Diseases (CIBERNED), Madrid, Spain.

Running title: Folate receptor role in rejuvenation of DG cells.

**Abstract**

In this work, we have studied the effect of small compounds in the partial rejuvenation of dentate gyrus cells by measuring the improvement of cognitive functions in elderly mice.
Aging has been related to a change in DNA methylation and some one-carbon metabolites linked to that methylation process, like vitamin B12, folate or methionine have been involved in cognitive performance during aging. However, their role in this process and the possible mechanisms behind its cognitive effects are still unclear. Through direct infusion of these molecules in the dentate gyrus we have tested their effects on cognition in elderly mice. Only positive results were found for folate. A partial rejuvenation of dentate gyrus cells related to an increase in neuroplasticity by reorganizing extracellular matrix structures and rising the expression of juvenile genes like GluN2B was found. Since folate is involved in several cellular pathways in addition to DNA methylation, we have focused in its interaction with its folate receptor alpha (FRα), a protein that is present at the cell nucleus, acting as transcription factor. We have found that most of folate effects on brain would be mediated by the activation of FRα. In addition, we propose that the mechanism for cell rejuvenation by folate, or other FRα binding molecules, may involve the expression of proteins, like SOX2, a Yamanaka factor present in young neurons. Thus, the use of molecules that activate the FRα pathway could constitute an interesting strategy to be considered for the study of brain rejuvenation.

**Keywords:** Aging; Folate; Rejuvenation; Cognition; Folate Receptor; Hippocampus

### Results

1. Effect of folate on adult neurogenesis in the dentate gyrus

As in our previous work [1], we tested the effect of folate on neuronal precursor cells, studying changes on brain lipid-binding protein (blbp) (**SI 4a**) and doublecortin-immunoreactive (ir) cell densities. Also, to label dividing cells, mice were injected intraperitoneally with CldU a week before perfusion, thus allowing the study of new-born 1-week-old cells in the SGZ. A single injection of folate into the hippocampus led to only a subtle increase in new-born CldU-ir cells (**SI 4b**) and in 1-week-old neurons (Dcx and CldU double-positive cells) (**SI 4c**) in the SGZ. Although the increase was not statistically significant, the tendency was similar to that found for YF [1]. Furthermore, we found no differences in Blbp-ir (**SI 4a**) or doublecortin-ir cell densities (**SI 4d**) upon folate injection. Therefore, folate had no significant effect on adult hippocampal neurogenesis (AHN).

Indeed, in our previous work [1], a major YF-dependent change was found only in developmentally generated neurons compared to neurons raised in adult neurogenesis in the hippocampal region. Thus, we subsequently tested the effect of folate on developmentally generated cells in the DG, looking at specific markers like changes in histone or DNA methylation.

2. Impact of folate on the methylation of histone and DNA in hippocampal neurons

The expression of aging-related genes correlates, as suggested by the heterochromatin loss aging model [2], with increasing alterations of chromatin located in heterochromatin regions. In this regard, we examined whether folate injection modifies the degree of histone methylation, namely at positions H3K9me3 and H4K20me3, which are present mainly in heterochromatin regions. The presence of folate 14 days post-injection did not modify the level of methylation at H3K9me3 (**SI 5a, b**) or H4K20me3 (**SI 5c, d**).

Our findings suggest that a single injection of folate might not be enough to promote reliable changes in histone methylation levels. Therefore, we tested whether folate affects DNA methylation. Significant increases in 5mC were observed in the granular neuronal layer (P=0.0132) of the DG where the injection was performed and it also tended to increase in the pyramidal layer of CA3 (P-value=0.05) and CA1 (P-value=0.06) (**SI 6d, e**)

Regarding 5hmC, we have also tested its levels, since the accumulation of 5hmC in postmitotic neurons is related to “demethylation” (5mC -> 5hmC) facilitating gene expression. No significant changes were found in hippocampal neuronal layers (**SI 7**).

### Legend to Figures

**Supplementary Figure 1. Experimental timeline and intracerebral injections. (a)** Schematic illustration of the experimental timeline of experiments performed in the present work. **(b)** Illustration (modified from Paxinos, George, and Keith B.J. Franklin. The mouse brain in stereotaxic coordinates: hardcover edition. Access Online via Elsevier, 2001) of anatomical localization of the bilateral monodose injections (red plot) performed in the Dentate Gyrus (Anteroposterior -2 mm; Mediolateral ±1.4 mm; Dorsoventral -2.2 mm) of different metabolites (vitamin B12, SAM, and folate) and of FRα-binding peptide or vehicle solution, in wild-type mice (male and female were randomized into experimental groups). On the bottom it is shown representative examples of trial hippocampal injections with dextran tracer (in red) performed before the start of the treatments. Note the trajectory of the 5ul Hamilton syringe, which only reaches the hilus of the DG.

**Supplementary Figure 2. Effect of vitamin B12 on behavior.** Single dose of hippocampal infusions of vitamin B12 did not alter the behavior of old wild-type mice compared with mice receiving vehicle solution. (**a**) Histograms show some of the main data from the open field test: time spent by mice in the center square of the box during the test, average speed of movement, and time spent immobile. (**b**) Histograms show results from two memory performance tests: the novel object recognition test (NOR) for short- and long-term recognition memory, and the Y maze test for spatial memory. Black bars represent mean± SD of the vehicle-treated group and gray bars represent mean± SD of the vitamin B12-treated group. All data were analyzed by Student's t-test.

**Supplementary Figure 3. Multiple folate effects, unique Frα binding peptide effects.** (**a**) As indicated in the text, folate may play a role in several different pathways. (**b**) However as far as we know the only role for Frα binding peptide (see next figures) could be to regulate the expression of specific genes. In that way Frα binding peptide could be a more suitable therapeutic agent for that gene expression regulation.

**Supplementary Figure 4. Effect of folate on the expression of adult neurogenesis markers**. (**a**) Histogram showing density of Blbp-ir cells per $mm^3$ obtained in the subgranular zone (SGZ) of the dentate gyrus. Single-dose treatment with folate did not cause statistically significant changes in early neural precursors in the SGZ. (**b**) Histogram showing density of CldU-ir cells per $mm^3$ in the SGZ. Single-dose treatment with folate did not cause statistically significant changes in the generation of new cells in the SGZ. (**c**) Histogram showing percentage of doublecortin-ir cells that were also CldU-immunoreactive in the SGZ. Single-dose treatment with folate did not cause statistically significant changes in the density of new 2-week-old neurons in the SGZ. (**d**) Histogram showing density of doublecortin-ir cells per $mm^3$ in the SGZ. Single-dose treatment with folate did not cause statistically significant changes in the density of young neurons in the SGZ. (Mean ± SD; ns: not significant differences; Student`s t test).

**Supplementary Figure 5. Levels of H3K9me3 and H4K20me3. Effect of folate**. (**a**) Representative stitched confocal images of H3K9me3 marker (in red) in the dentate gyrus (DG). Inset also shows the DAPI signal (in blue) to identify the DG structure. (**b**) Histogram showing mean intensity of histone methylation signal. The effect of the absence (black) or presence (gray) of folic acid is shown. (**c**) Representative stitched confocal images of the H4K20me3 marker (in green) in the DG. Inset also shows the DAPI signal (in blue) to identify the DG structure. (**d**) Histogram showing mean intensity of histone methylation signal. The effect of the absence (black) or presence (gray) of folic acid is shown. (Mean ± SD; ns: not significant differences; Student`s t test). Scale bar, 100 µm.

**Supplementary Figure 6. Effect of folate on DNA methylation of hippocampal neurons.** Representative stitched confocal images of 5-methycytosine (5mC) marker (in green) in dentate gyrus of vehicle-treated (**a**) or folate treated (**b**). the corresponding high magnifications of the granular cell layer of a and b (yellow squares) are shown. Heterochromatin clusters of 5mC immunoreactivity can be observed in a' and b'. (**c**), (**d**) and (**e**) histograms showing mean intensity of the 5mC signal in the granular cell layer CA3 and CA1 hippocampal regions. The effect of the absence (black) or presence (gray) of folic acid is shown (mean ± SD, *$p<0.05$. Student's test; ns: not significant differences).

**Supplementary Figure 7. Levels of 5hmC. Effect of folate**. (**a**) Representative stitched confocal images of 5-hydroxymethylcytosine (5hmC) marker (in red) in the dentate gyrus (DG) of vehicle-treated and folate-treated mice. (**b**) Histogram showing mean intensity from 5mC signal of the granular cell layer of the DG. The effect of the absence (black) or presence (gray) of folic acid is shown. (Mean ± SD; *$p<0.05$, Student`s t test; ns: not significant differences). Scale bar, 100 µm.

**Supplementary Figure 8. Expression of the FRα-binding peptide on nuclear fraction of SK-N-SH cells by addition of folate or the FRα-binding peptide.** Boshnjaku et al. [3]described that, upon folate addition, FRα-binding peptide (which appears as two bands around 42kD and 38kD) can be internalized to the cytoplasm (~42 kD band) or the nucleus (~38kD band). We therefore studied the nuclear localization of FRα-binding peptide upon addition to SK-N-SH human neuroblastoma cells of two concentrations of folate and FRα-binding peptide (0.5 (+) and 1 (++) mM). Our results show the presence of ~38Kd band in the nuclear fraction (**a**) in the presence or absence of folate or FRα-binding peptide at different concentrations. (**b**) Quantification of the results shown in a. Results of densitometry were normalized against the control value. (**c**) Folate or FRα-binding peptide addition results in an increase in the expression of Sox2 protein. (**d**) Quantification of the results shown in c.

**Supplementary Figure 9. Levels of 5mC and 5hmC. Effect of the FRα-binding peptide**. Histograms showing mean intensity from 5mC (**a**) and 5hmC (**b**) signal of the granular cell layer of the dentate gyrus (DG) region. The effect of the absence (black) or presence (gray) of the FRα-binding peptide (2,5mg/ml) is shown. No significant effect on DNA methylation status of the DG was found. (Mean ± SD; $^*p<0.05$, Student`s t test; ns: no significant differences).


1. Rodriguez-Matellan, A., et al., *In Vivo Reprogramming Ameliorates Aging Features in Dentate Gyrus Cells and Improves Memory in Mice.* Stem Cell Reports, 2020. **15**(5): p. 1056-1066.
2. Villeponteau, B., *The heterochromatin loss model of aging.* Exp Gerontol, 1997. **32**(4-5): p. 383-94.
3. Boshnjaku, V., et al., *Nuclear localization of folate receptor alpha: a new role as a transcription factor.* Sci Rep, 2012. **2**: p. 980.

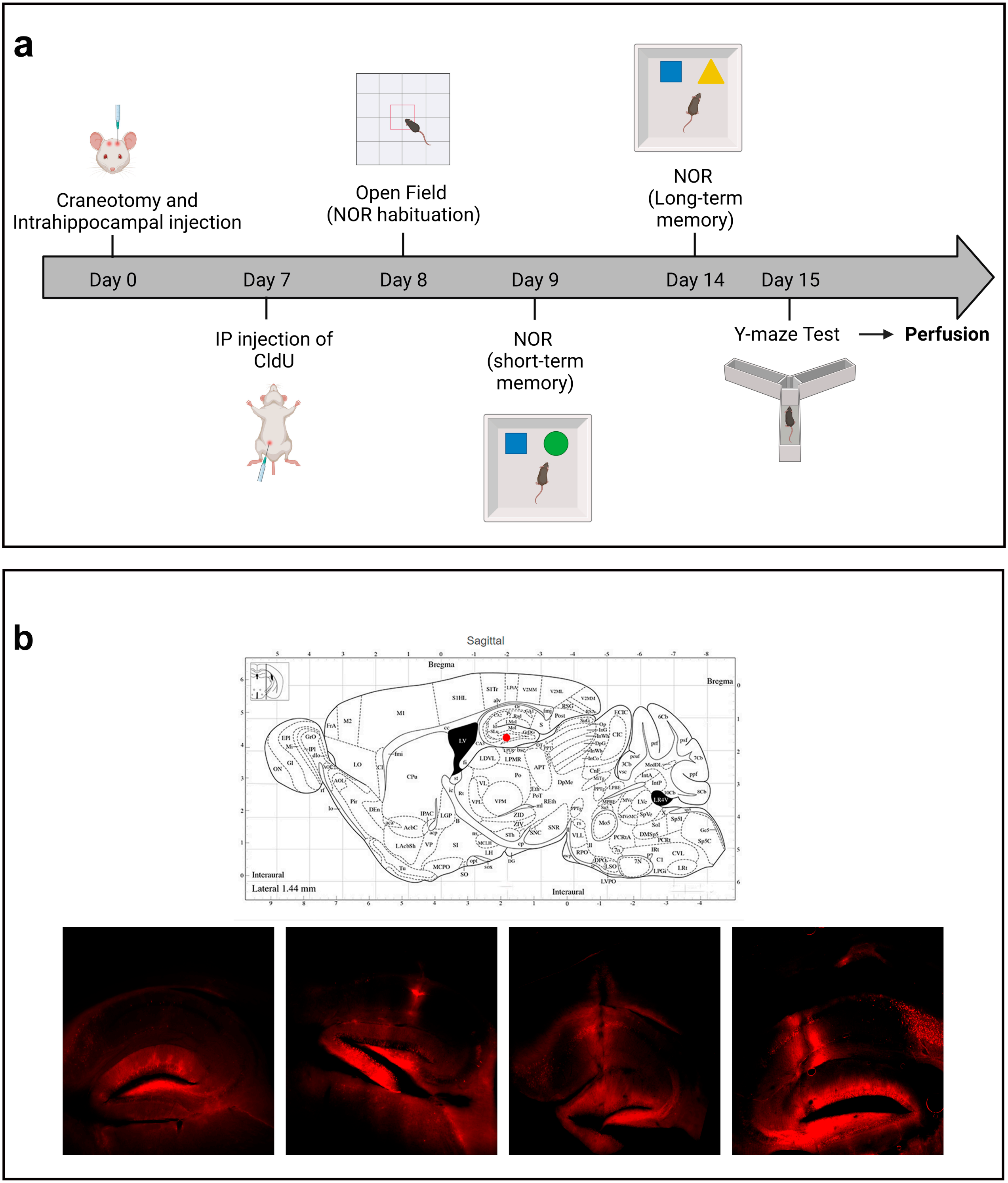
a
Craneotomy and
Intrahippocampal injection
Open Field
(NOR habituation)
NOR
(Long-term
memory)
Day 0
Day 7
Day 8
Day 9
Day 14
Day 15
IP injection of
CldU
NOR
(short-term
memory)
Y-maze Test
Perfusion
b
Sagittal
Bregma
Interaural
Lateral 1.44 mm

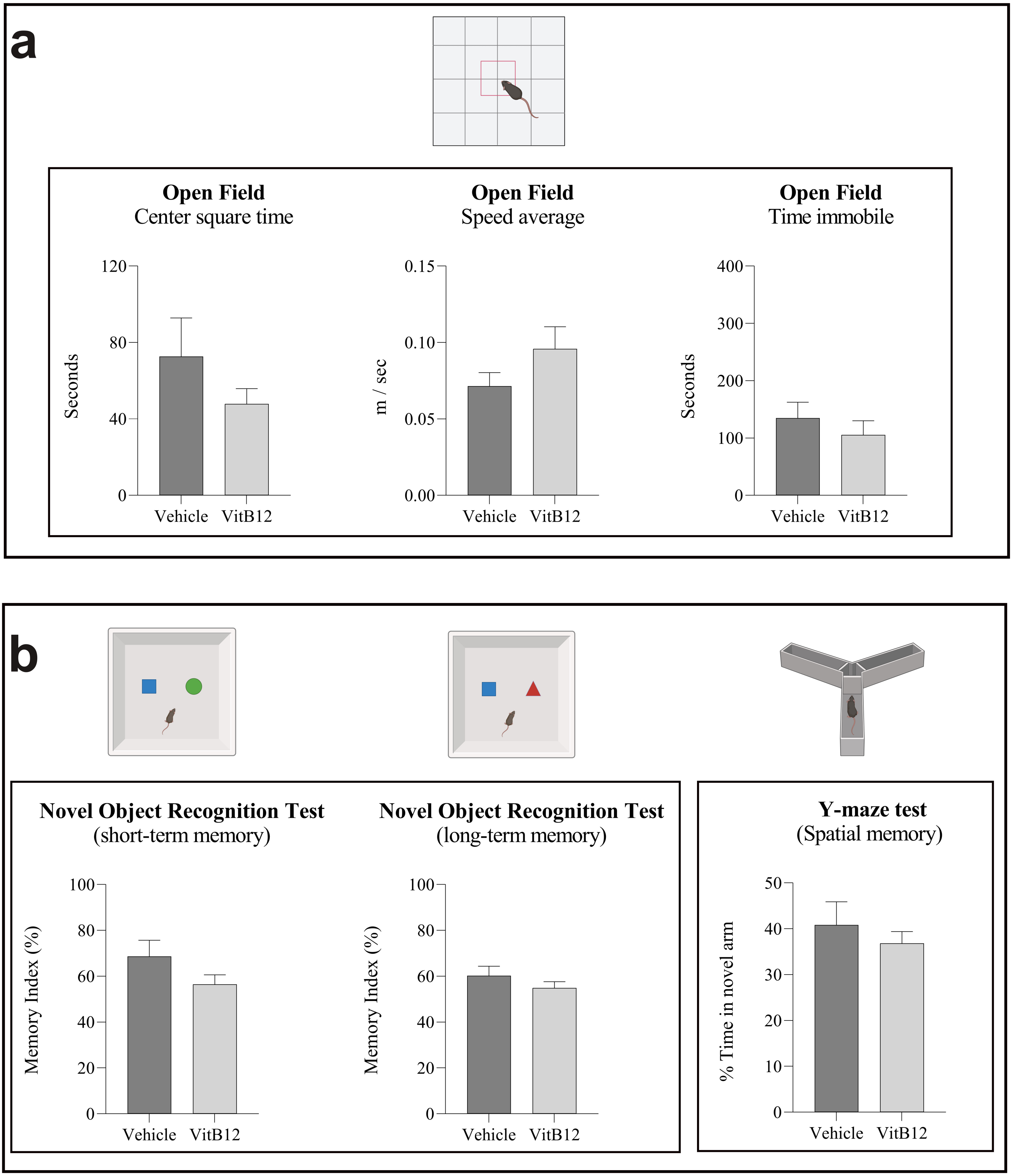
a
Open Field
Center square time
120
80
40
0
Seconds
Vehicle
VitB12
Open Field
Speed average
0.15
0.10
0.05
0.00
m / sec
Vehicle
VitB12
Open Field
Time immobile
400
300
200
100
0
Seconds
Vehicle
VitB12
b
Novel Object Recognition Test
(short-term memory)
100
80
60
40
20
0
Memory Index (%)
Vehicle
VitB12
Novel Object Recognition Test
(long-term memory)
100
80
60
40
20
0
Memory Index (%)
Vehicle
VitB12
Y-maze test
(Spatial memory)
50
40
30
20
10
0
% Time in novel arm
Vehicle
VitB12

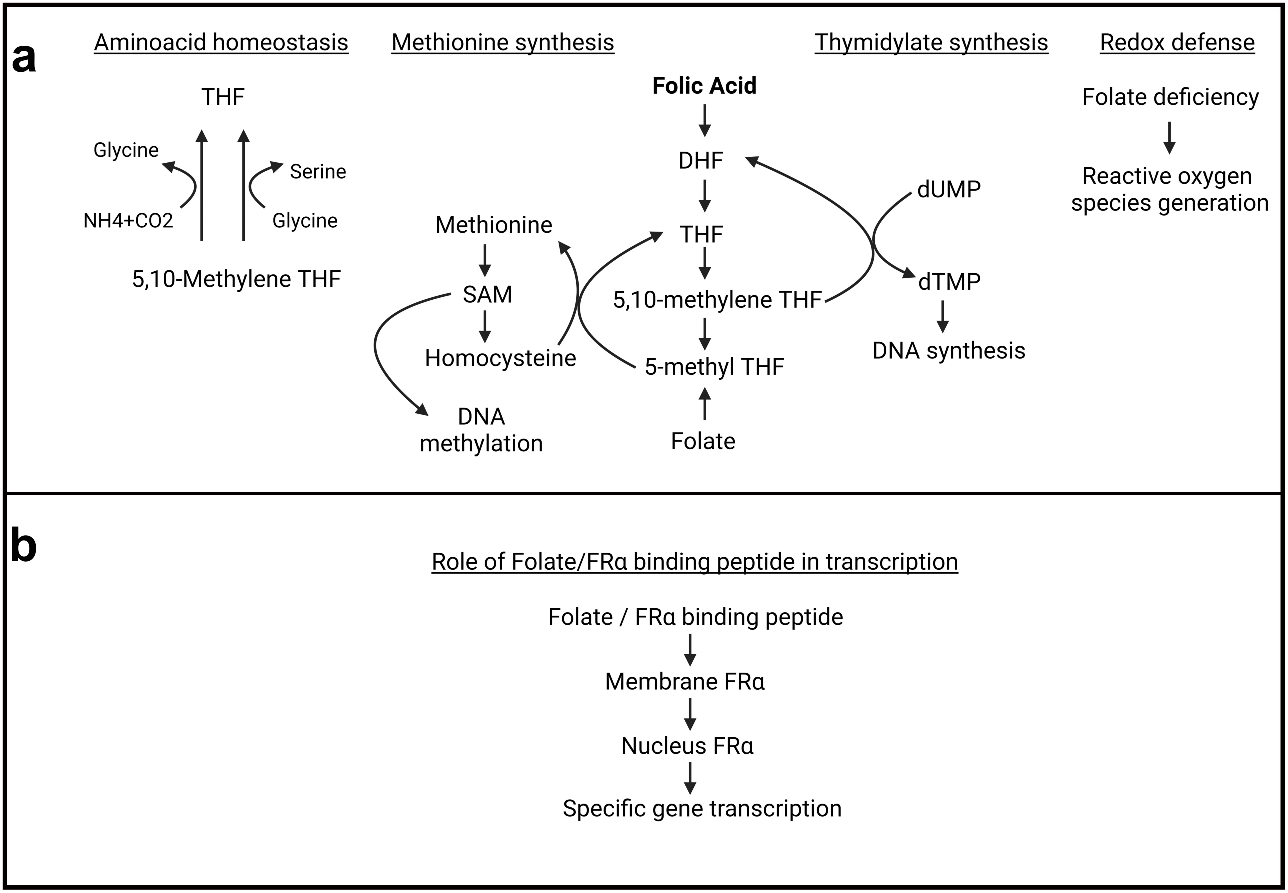
a
Aminoacid homeostasis
Methionine synthesis
Thymidylate synthesis
Redox defense
THF
Glycine
Serine
NH4+CO2
Glycine
5,10-Methylene THF
Folic Acid
DHF
THF
5,10-methylene THF
5-methyl THF
Folate
Methionine
SAM
Homocysteine
DNA methylation
dUMP
dTMP
DNA synthesis
Folate deficiency
Reactive oxygen species generation
b
Role of Folate/FRα binding peptide in transcription
Folate / FRα binding peptide
Membrane FRα
Nucleus FRα
Specific gene transcription

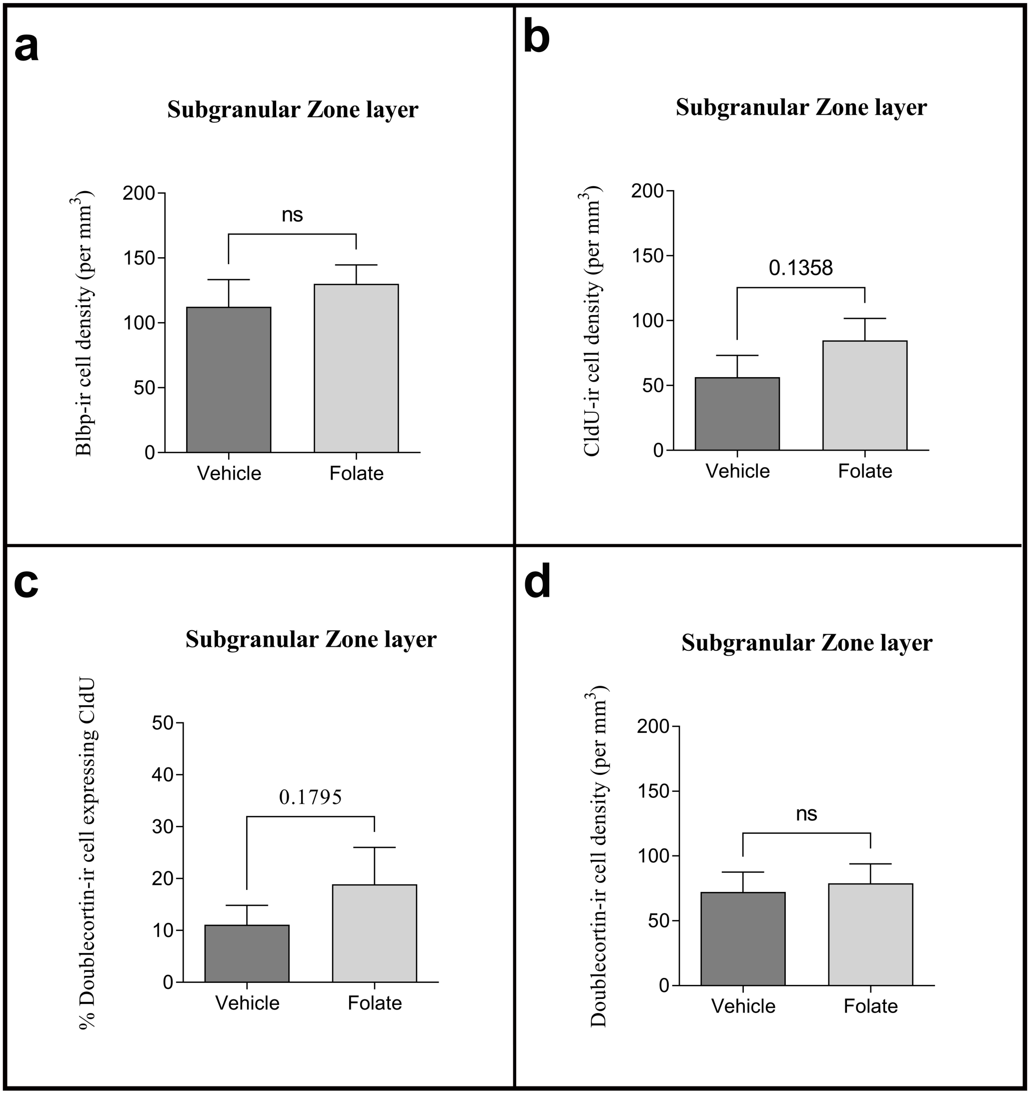
a
Subgranular Zone layer
Blbp-ir cell density (per mm3)
200
150
100
50
0
ns
Vehicle
Folate
b
Subgranular Zone layer
CldU-ir cell density (per mm3)
200
150
100
50
0
0.1358
Vehicle
Folate
c
Subgranular Zone layer
% Doublecortin-ir cell expressing CldU
50
40
30
20
10
0
0.1795
Vehicle
Folate
d
Subgranular Zone layer
Doublecortin-ir cell density (per mm3)
200
150
100
50
0
ns
Vehicle
Folate

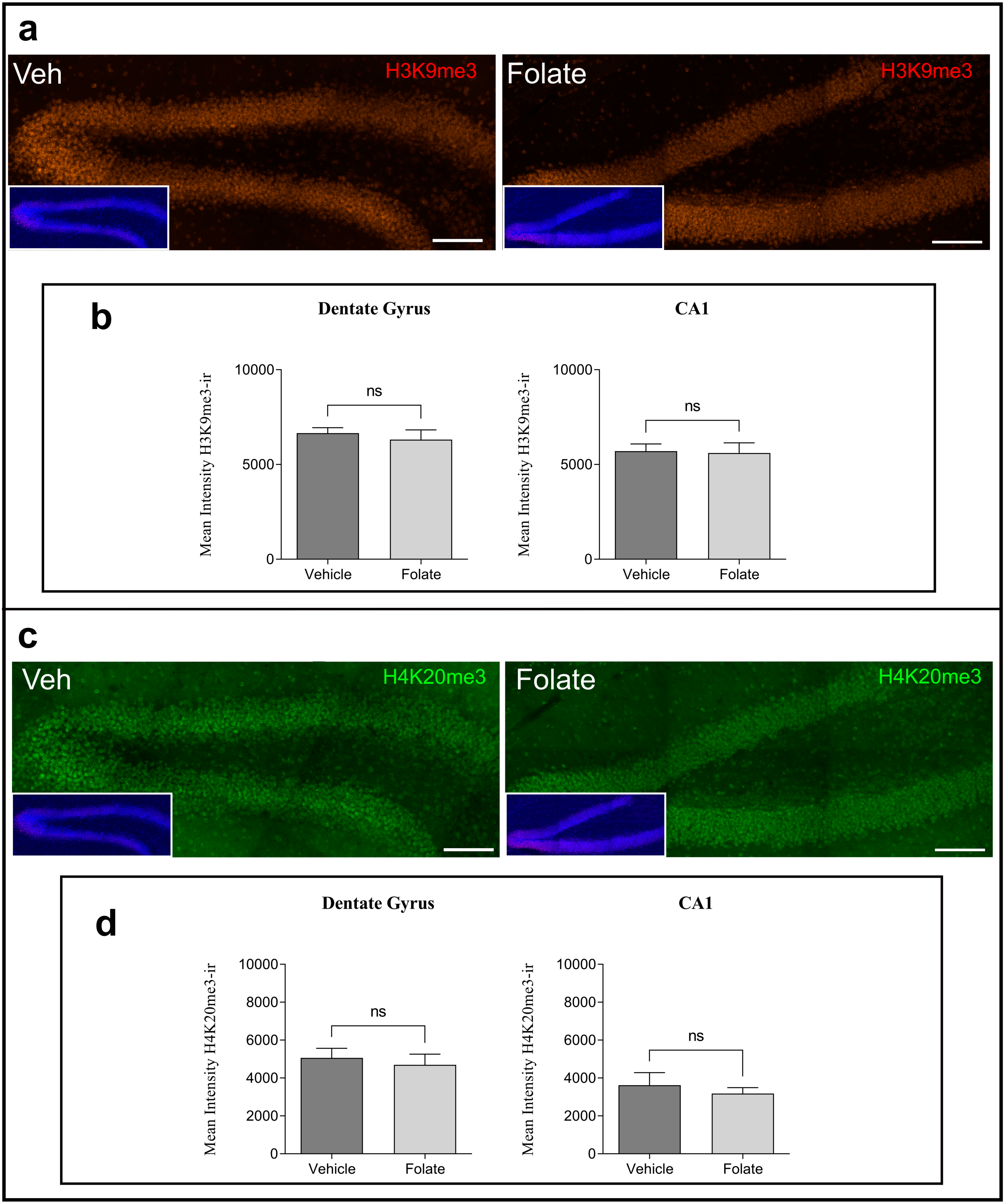
a
Veh
H3K9me3
Folate
H3K9me3
b
Dentate Gyrus
CA1
Mean Intensity H3K9me3-ir
10000
5000
0
ns
Vehicle
Folate
Mean Intensity H3K9me3-ir
10000
5000
0
ns
Vehicle
Folate
c
Veh
H4K20me3
Folate
H4K20me3
d
Dentate Gyrus
CA1
Mean Intensity H4K20me3-ir
10000
8000
6000
4000
2000
0
ns
Vehicle
Folate
Mean Intensity H4K20me3-ir
10000
8000
6000
4000
2000
0
ns
Vehicle
Folate

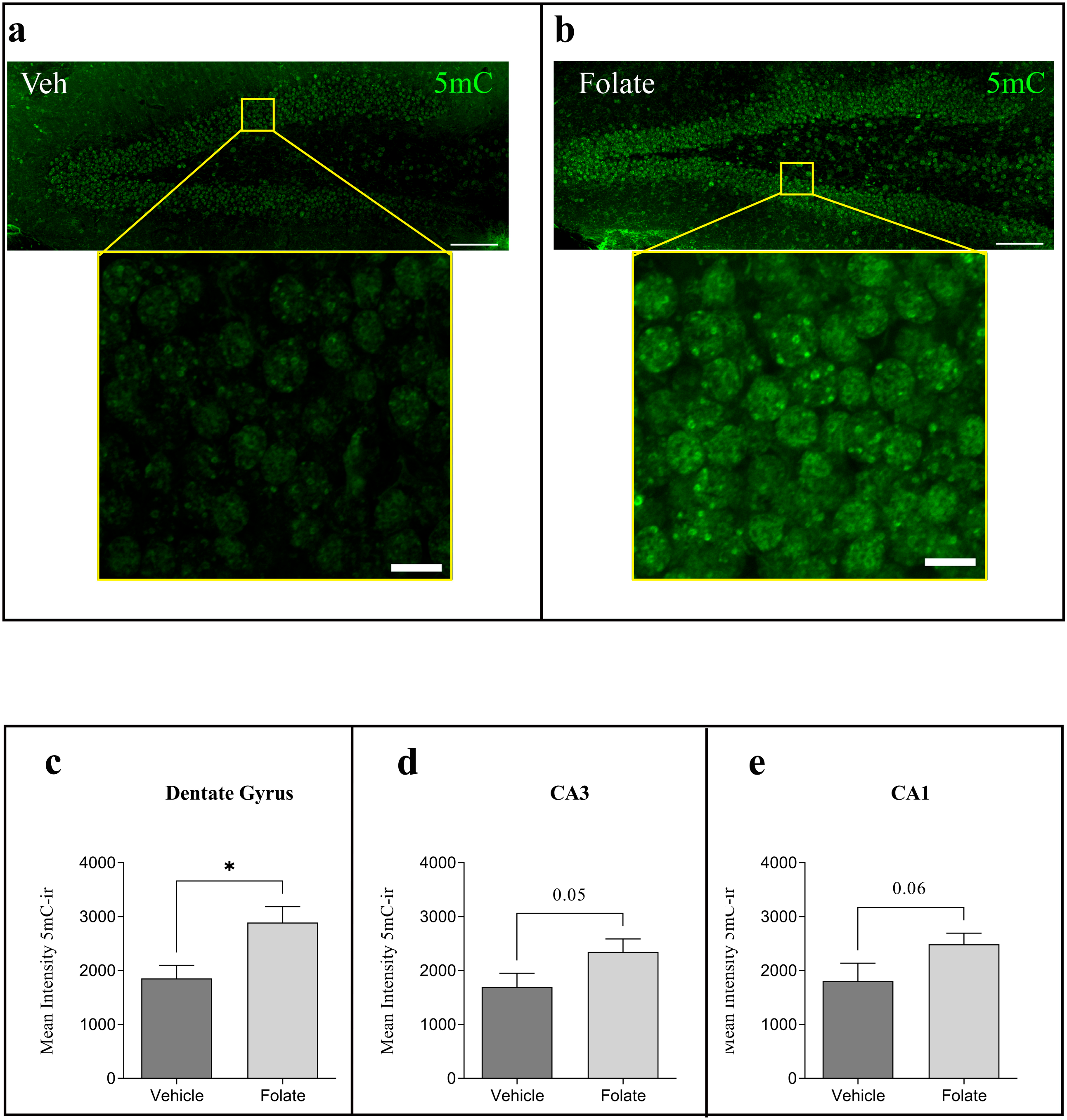

a
Veh
5mC
b
Folate
5mC
c
Dentate Gyrus
Mean Intensity 5mC-ir
4000
3000
2000
1000
0
*
Vehicle
Folate
d
CA3
Mean Intensity 5mC-ir
4000
3000
2000
1000
0
0.05
Vehicle
Folate
e
CA1
4000
3000
2000
1000
0
0.06
Vehicle
Folate

a

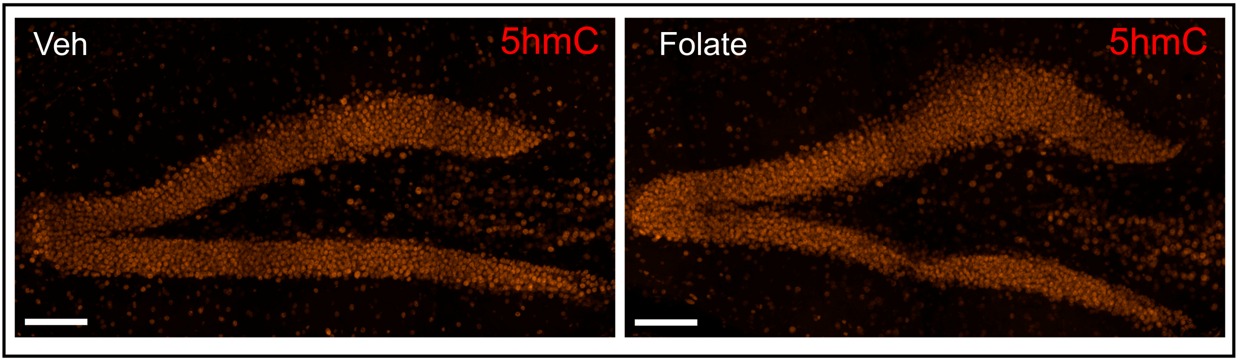
Veh
5hmC
Folate
5hmC

b

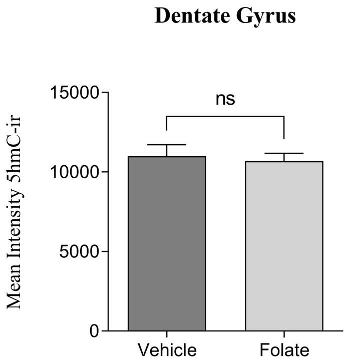
Dentate Gyrus
ns
15000
10000
5000
0
Mean Intensity 5hmC-ir
Vehicle
Folate

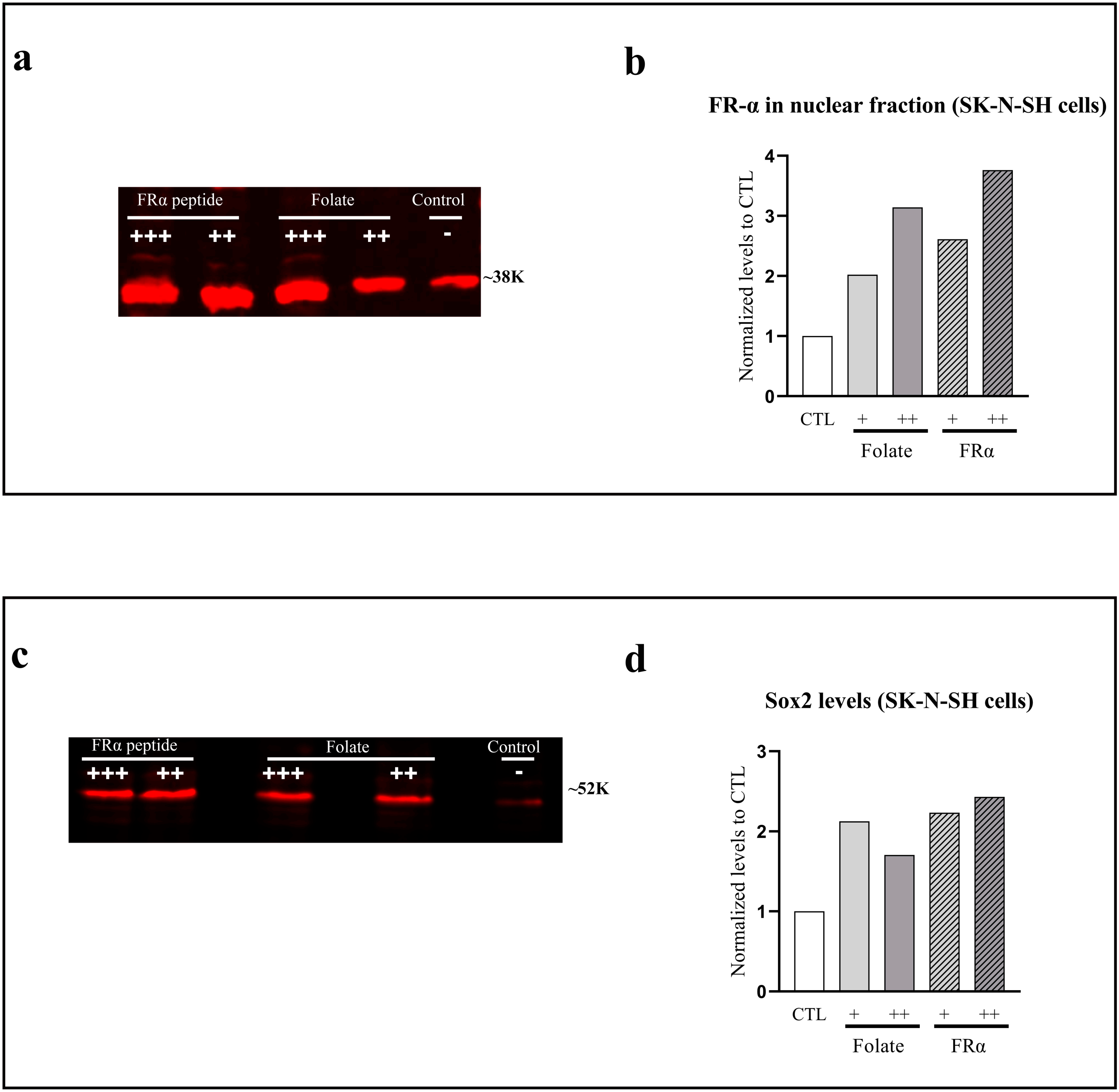
a
FRα peptide
Folate
Control
+++
++
+++
++
-
~38K
b
FR-α in nuclear fraction (SK-N-SH cells)
Normalized levels to CTL
0
1
2
3
4
CTL
+
++
+
++
Folate
FRα
c
FRα peptide
Folate
Control
+++
++
+++
++
-
~52K
d
Sox2 levels (SK-N-SH cells)
Normalized levels to CTL
0
1
2
3
CTL
+
++
+
++
Folate
FRα

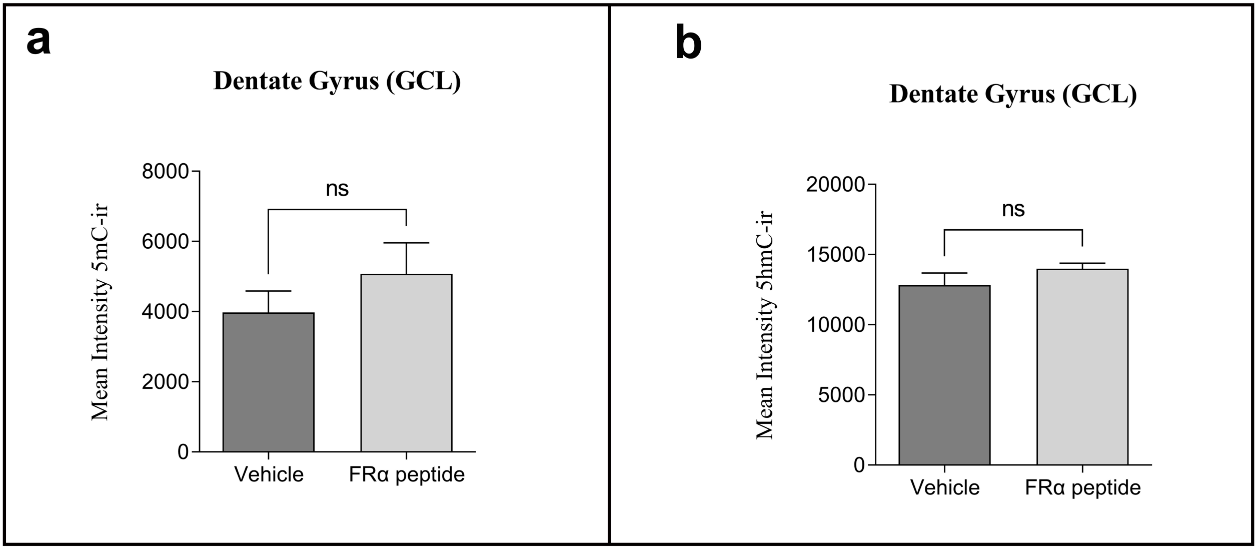
a
Dentate Gyrus (GCL)
Mean Intensity 5mC-ir
8000
6000
4000
2000
0
ns
Vehicle
FRα peptide
b
Dentate Gyrus (GCL)
Mean Intensity 5hmC-ir
20000
15000
10000
5000
0
ns
Vehicle
FRα peptide
